# Quantitative contributions of hepatic and renal organic cation transporters to the clinical pharmacokinetic cimetidine-metformin interaction

**DOI:** 10.1101/2024.11.19.624371

**Authors:** Anoud Sameer Ailabouni, Dilip Kumar Singh, Aarzoo Thakur, Mary F. Paine, Erin C. Boone, Andrea Gaedigk, Bhagwat Prasad

**Affiliations:** College of Pharmacy and Pharmaceutical Sciences, Washington State University, Spokane; Children’s Mercy Research Institute, Division of Clinical Pharmacology, Toxicology & Therapeutic Innovation, Kansas City; University of Missouri-Kansas City, School of Medicine, Department of Pediatrics, Kansas City

**Keywords:** Biomarkers, drug-drug interactions, hepatic, metformin, organic cation transporters, pharmacogenetics, pharmacokinetics, physiology-based pharmacokinetic, renal

## Abstract

The widely prescribed oral anti-diabetic drug metformin is eliminated unchanged in the urine primarily through active tubular secretion. This process is mediated by organic cation transporter 2 (OCT2), an uptake transporter expressed on the basolateral membrane of renal proximal tubule cells. Metformin uptake into the liver, the site of action, is mediated by OCT1, which is expressed on the sinusoidal membrane of hepatocytes. Sixteen healthy adults participated in a clinical pharmacokinetic drug-drug interaction study in which they were orally administered metformin (50 mg) as a dual OCT1/2 substrate alone (baseline) and with cimetidine (400 mg) as an OCT inhibitor. Relative to baseline, metformin systemic plasma exposure increased by 24% (*p*<0.05) in the presence of cimetidine, which was accompanied by a disproportional decrease (8%) in metformin renal clearance (*p*=0.005). Genetic variants of *OCT1* and *OCT2* moderately impacted the significance and magnitude of the interaction. Collectively, we hypothesized that the cimetidine-metformin interaction involves inhibition of hepatic OCT1 as well as renal OCT2. We tested this hypothesis by developing a physiologically based pharmacokinetic (PBPK) model and assessing potential OCT biomarkers in plasma and urine to gain mechanistic insight into the transporters involved in this interaction. The PBPK model predicted that cimetidine primarily inhibits hepatic OCT1 and, to a lesser extent, renal OCT2. The unchanged renal clearance of potential OCT2 biomarkers following cimetidine exposure supports a minimal role for renal OCT2 in this interaction.

## INTRODUCTION

Metformin is a first-line treatment for type 2 diabetes mellitus, ranking among the most widely prescribed medications globally.^1^ The primary mechanism of action of metformin involves phosphorylation of 5’-adenosine monophosphate kinase (AMPK), resulting in inhibition of excessive hepatic glucose production by reducing gluconeogenesis.^2^ Metformin, a biguanide, is highly hydrophilic (log P, -1.43) and carries a positive charge at physiological pH due to its high pKa (11.5),^3^ requiring cationic transporters for uptake and efflux through cell membranes.

Metformin uptake into the liver, the major site of action, is primarily mediated by organic cation transporter 1 (OCT1), which is expressed on the basolateral membrane of hepatocytes.^1^ In *Oct1* knockout mice, hepatic metformin concentration was significantly reduced by 3.4-fold compared to that in wild-type mice after a single oral dose of metformin, resulting in a reduction in AMPK phosphorylation and glucose-lowering effect.^4^ These observations suggest that Oct1/OCT1 activity plays a crucial role in facilitating hepatic metformin uptake and subsequent pharmacological response. Metformin is not metabolized and is excreted unchanged into the urine, primarily through renal secretion (75%), along with glomerular filtration (25%).^5^ The population mean plasma renal clearance (CL_r_) of metformin in healthy adults is 510±120 mL/min, representing a substantive fraction of the effective renal plasma flow (∼600 mL/min).^6^ Metformin uptake into the kidney is facilitated by OCT2, which is predominantly expressed on the basolateral membrane of renal proximal tubule cells.^1^ Renal secretion of metformin into the urine is mediated in conjunction with multidrug and toxic extrusion 1 (MATE1) and MATE2K, which are efflux transporters expressed on the apical membrane of renal proximal tubule cells.^1^

Genetic variation in the genes coding OCT1 (*SLC22A1),* OCT2 (*SLC22A2)*, MATE1 (*SLC47A1)* and/or MATE2K (*SLC47A2*) can impact metformin pharmacokinetics and pharmacodynamics. For simplicity, we are referring to the genes as *OCT1, OCT2, MATE1* and *MATE2K* throughout this report. The pharmacokinetic profiles of a single oral dose of metformin (500 mg) in individuals with *OCT2* c.596C>T (p.Thr199Ile, rs201919874), c.602C>T (p.Thr201Met, rs145450955), and/or c.808G>T (p.Ser270Ala, rs316019) were associated with a decrease in transporter function compared to individuals with the reference genotype. These variants demonstrated significantly higher plasma exposure and reduced CL_r_ of metformin compared with the reference group.^7^ Additionally, the *OCT1* intronic variant c.1386-2964C>A (rs622342), which is associated with altered OCT1 function, resulted in an impaired glucose-lowering effect.^8^

Administration of OCT/MATE inhibitors (e.g., cimetidine) can alter metformin pharmacokinetics and pharmacodynamics. Pharmacokinetic cimetidine-metformin interactions have been reported in both humans and rodents. For example, coadministration of cimetidine (400 mg twice daily for 6 days) to healthy adults increased steady state plasma concentrations of metformin (250 mg) by approximately 50% and decreased CL_r_ by 27%, suggesting cimetidine inhibited OCT2 and/or MATE1/2K.^9^ Likewise, we recently showed that coadministration of cimetidine (100 mg/kg, intraperitoneally) to rats significantly increased metformin blood concentrations (3.2-fold) and reduced CL_r_ (73%).^10^ Consistent with these results, cimetidine was shown to significantly increase the liver-to-plasma ratio of metformin in mice.^11^ The contribution of OCT1 to the cimetidine-metformin interaction in humans remains unclear.

Endogenous transporter substrates have emerged as potential tools to predict transporter-mediated drug-drug interactions (DDIs).^12^ Recent studies proposed N1-methylnicotinamide (NMN) and N1-methyladenosine (NMA) as potential biomarkers of OCT2. For example, following administration of cimetidine (400 mg) 1 hour before metformin (500 mg) to healthy adults, NMN CL_r_ decreased by approximately 35%, suggesting NMN could be used as a renal cation transporter biomarker.^13^ Similarly, CL_r_ of NMA was reduced by 10%, 21%, and 47% following administration of 10, 25, and 75 mg of pyrimethamine, respectively, to healthy adults, indicating renal cation transporter inhibition.^14^ Regarding OCT1, isobutyryl-L-carnitine (IBC) has been proposed as a potential endogenous substrate. A first-in-human study showed a potent OCT1 inhibitor new chemical entity to significantly decrease plasma exposure to IBC by 35% in healthy adults compared to placebo.^15^ Although NMN, NMA, and IBC have been proposed as potential OCT1 and/or OCT2 biomarkers, their applicability as sensitive and selective biomarkers for these transporters remain uncertain.^12^

Despite several studies indicating roles for OCTs in metformin pharmacokinetics, individual contributions of OCT1 versus OCT2 remain unknown. Because cimetidine is a dual OCT1/2 inhibitor, we hypothesized that cimetidine inhibits both hepatic OCT1 and renal OCT2, resulting in a disproportional increase in plasma concentrations versus a decrease in the CL_r_ of metformin (Figure 1) and OCT biomarkers (i.e., NMN, NMA, and IBC). We tested this hypothesis using an integrated, three-pronged approach: 1) conduct a pharmacokinetic cimetidine-metformin interaction study in healthy adults to quantify plasma concentrations and determine CL_r_ of metformin and OCT biomarkers in the absence and presence of cimetidine, 2) develop a mechanistic physiologically based pharmacokinetic (PBPK) model to capture the contributions of hepatic OCT1 and renal OCT2 inhibition to the interaction, and 3) explore the impact of *OCT1*, *OCT2*, *MATE1* and *MATE2K* genetic variation on metformin pharmacokinetics and interaction with cimetidine. Our data suggests that hepatic OCT1 contributes to altered metformin DDI with cimetidine besides renal OCT2, which should be considered as a key mechanism of disposition of metformin while interpreting DDI and pharmacogenomic data.

**Figure 1.**
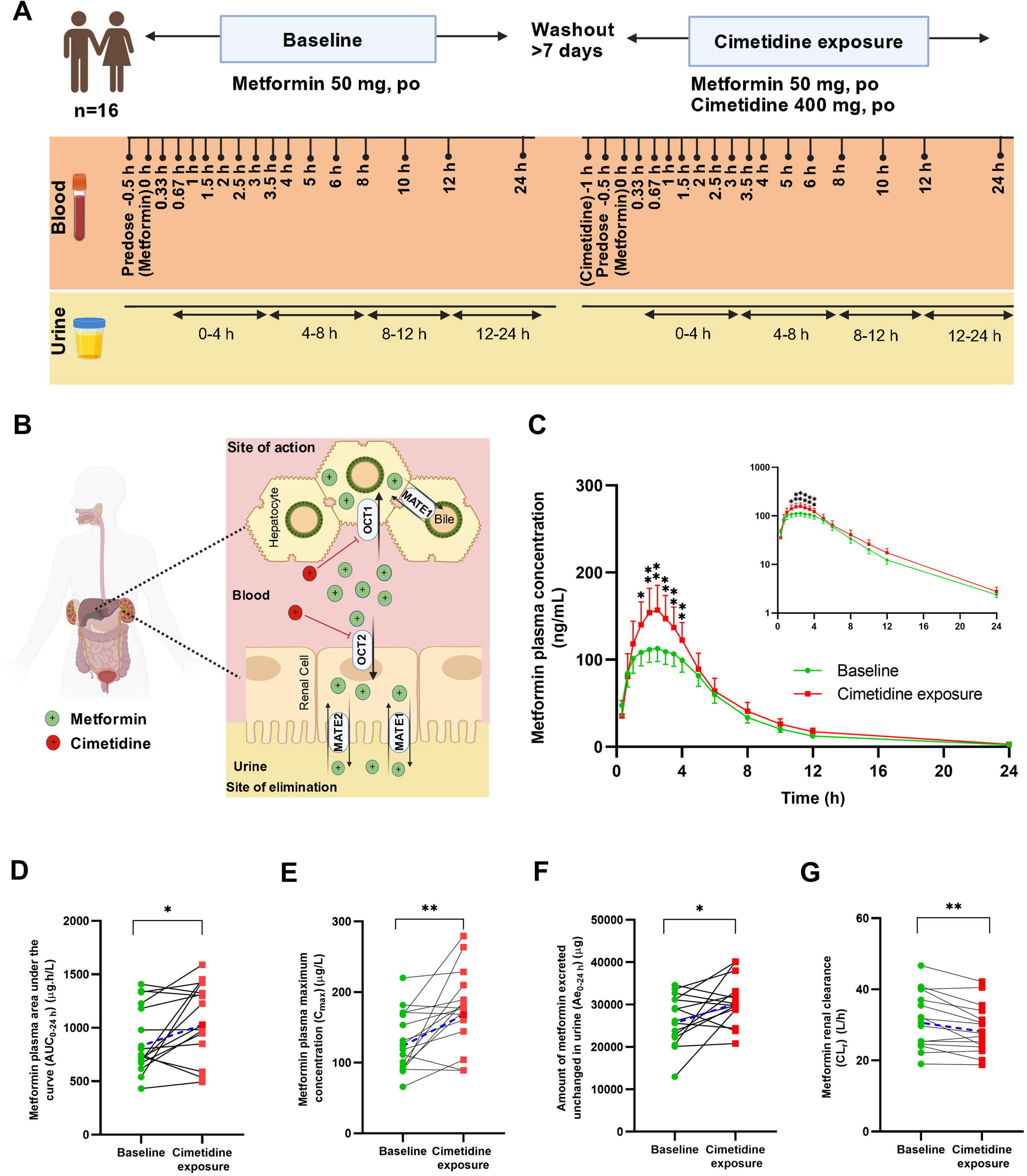
Effect of cimetidine co-administration on metformin pharmacokinetics. Clinical study design and dosing schedule (A). A schematic representation of transport processes involving metformin and its interaction with cimetidine in hepatocytes and renal proximal tubule cells in humans (B). Metformin pharmacokinetic measures, i.e., the mean plasma concentration-time profile (C), plasma area under the curve from 0-24 h (AUC_0-24 h_) (D), plasma maximum concentration (C_max_) (E), amount excreted unchanged into the urine from 0-24 h (Ae_0-24 h_) (F), and renal clearance (CL_r_) (G) of metformin when administered alone (baseline, green circles) and in combination with cimetidine (cimetidine exposure, red squares) in healthy adults (n=16). One participant was excluded from the Ae and CL_r_ analysis due to difficulty voiding urine. In Figure 1A, Metformin (green circles) is transported into hepatocytes via sinusoidal organic cation transporter 1 (OCT1) and secreted into urine through organic cation transporter 2 (OCT2) and multidrug and toxin extrusion (MATE) transporters expressed in the renal tubular cells. Cimetidine (red circles) inhibits hepatic OCT1 and renal OCT2. In Figure 1C, Symbols and error bars represent observed geometric means and 90% confidence intervals, respectively. In Figure 1G, CL_r_ was calculated based on (0-24 h) time interval. In Figure 1D-G, solid black lines indicate individual values and dashed line represents geometric mean. Statistical analysis was conducted using two-tailed paired t-tests: *p-value* < 0.05 (*) and < 0.01 (**). Figures 1A and B were created using BioRender.

## MATERIALS AND METHODS

### Materials

Metformin hydrochloride, cimetidine, metformin-d6, NMN-d3, and NMA-d3 were purchased from Toronto Research Chemicals (Toronto, Ontario, Canada). LC-MS grade methanol, acetonitrile, water, and formic acid were purchased from Fisher Scientific (Fair Lawn, NJ). Bovine serum albumin (BSA) was procured from Sigma-Aldrich (St. Louis, MO). Reagent grade (≥95 % purity) of NMN, NMA, and IBC were purchased from Cayman Chemical (Ann Arbor, MI).

### Clinical study participants

The Washington State University (WSU) Institutional Review Board approved the clinical protocol and informed consent form [clinical trial registry number, NCT05365451]. Sixteen healthy adults (eight males and eight nonpregnant, nonlactating females) aged 24-64 years and meeting the inclusion/exclusion criteria (Table S1) were enrolled in the study. Written informed consent was obtained from all participants before screening. They underwent a medical history review, physical exam, and blood and urine collection to obtain complete blood count with differential, electrolytes, liver function tests, serum creatinine, and routine urine analysis including a pregnancy test for females. None of the participants showed evidence of renal and hepatic dysfunction. Their demographic characteristics are provided (Table S2).

### Clinical study design

The pharmacokinetic cimetidine-metformin interaction study consisted of two arms, which were separated by at least one week (Figure 1A). Participants fasted overnight before each study day. During Arm 1 (baseline), they were administered a single oral subtherapeutic dose (50 mg) of metformin (500 mg/5 ml solution, Ranbaxy Pharmaceuticals, Jacksonville, FL). During Arm 2 (cimetidine exposure), they were administered a single oral dose (400 mg) of cimetidine (200-mg film-coated tablets, Mylan Pharmaceuticals, Inc., Morgantown, WV) one hour before administration of metformin. Participants continued to fast until 4 hours after metformin administration, when they were provided lunch. During both Arms, blood (5 mL) was collected from an indwelling venous catheter (placed in an antecubital vein) 0.5 h before and at 0.33, 0.67, 1, 1.5, 2, 2.5, 3, 3.4, 4, 5, 6, 8, 10, and 12 h after metformin administration. Blood was centrifuged immediately to harvest plasma. An additional blood sample was collected for genotyping purposes during Arm 1 as described below. Urine was collected into one or more containers at 0-4, 4-8, and 8-12 h intervals after metformin administration. After the last blood draw, participants were discharged, then returned the next morning for a single venous blood draw at 24 h. Upon discharge, they were instructed to collect urine until returning for their 12-24 h collection the next morning. Urine volume for each time interval was measured and recorded. Plasma and urine samples were stored at -80°C until analysis by liquid chromatography-tandem mass spectrometry (LC-MS/MS).

The bioanalytical method for OCT substrate analysis in plasma and urine is provided in the Supplementary File. LC parameters and gradient conditions and multiple reaction monitoring (MRM) transitions used to analyze metformin, cimetidine, NMN, NMA, IBC, other potential OCT substrates, calibration curve standards, and quality control (QC) samples are described in Tables S3 and S4, respectively. LC-MS/MS peak integration and quantification were performed using Skyline software 23.1.0.268. (University of Washington, Seattle, WA).

### Whole exome sequencing

Whole blood and residual blood mixture were prepared for DNA isolation as described in Supplementary File. Whole exome sequencing (WES) was performed by the Children’s Mercy Genomics Core Service Center (Kansas City, MO). To investigate the association of genetic variation in *OCT1, OCT2, MATE1*, and *MATE2K*, with metformin pharmacokinetics and/or cimetidine-metformin DDI, exonic and selected intronic single nucleotide polymorphisms (SNPs) found in three or more participants were included for analysis. For each variant, participants were categorized into one of three groups: reference allele, heterozygous alternative (variant) allele, or homozygous alternative (variant) allele. If either the heterozygous or homozygous group had fewer than three participants, they were combined into a single alternative (variant) allele group to enable statistical analysis. SNPs found in fewer than three participants assigned to either reference or alternative (combined heterozygous or homozygous) alleles were excluded as statistical analysis was not applicable.

### Pharmacokinetic analysis

Area under the plasma concentration-time curve from 0 to 24 h (AUC_(0-24 h)_) and plasma maximum concentration (C_max_) were determined using non-compartmental analysis methods and MATLAB^®^ (R2023b; Natick, MA). Analyte concentrations in urine from 0 to 24 h were converted to the amount excreted (A_e(0-24 h)_) by multiplying with urine volume. Pharmacokinetic measures were determined based on 0-4 h time intervals, (AUC_(0-4 h)_ and A_e(0-4 h)_) during which the highest cimetidine plasma concentrations were observed. Renal clearance (CL_r_) was calculated using two methods: CL_r,a_ by dividing A_e(0-24 h)_ by AUC_(0-24 h)_ and CL_r,b_ by dividing A_e(0-4 h)_ by AUC_(0-4 h)_.

Percent changes in AUC, C_max_, A_e_, and CL_r_ in the presence relative to the absence of cimetidine were calculated to evaluate the DDI. Percent changes in AUC_(0-24 h)_ and C_max_ in the presence relative to the absence of cimetidine were calculated to evaluate the effects of *OCT1* genetic variants on the interaction. Percent changes in AUC_(0-24 h)_ and CL_r,a_ in the presence relative to the absence of cimetidine were calculated to evaluate the effects of *OCT2*, *MATE1*, and *MATE2K* genetic variants on the interaction. Detailed statistical analysis is provided in the Supplementary File.

### PBPK model development and verification of the cimetidine-metformin interaction

A PBPK model was developed to investigate the DDI between metformin and cimetidine using Simcyp software (version 22.0, Certara, Sheffield, UK) to gain mechanistic insights into the contributions of hepatic OCT1 and renal OCT2 to the interaction. System-dependent parameters were automatically incorporated from the software. Likewise, most drug-dependent parameters for metformin and cimetidine were incorporated into the software library with adjustments made to some parameters related to metformin and cimetidine transport kinetics to align with our observed data. Metformin- and cimetidine-specific input parameters are provided in Tables S5 and S6. Detailed model development and verification are explained in Supplementary File.

## RESULTS

### Effects of cimetidine co-administration on metformin pharmacokinetics

Both study drugs were well-tolerated by all participants, with minimal to no adverse effects. Blood samples were collected from all participants during both arms at all pre-defined times. Urine samples were collected from all participants during both arms except for one participant during the cimetidine exposure arm who had difficulty voiding urine. A validated bioanalytical method (Supplementary File) was used to quantify cimetidine, metformin, and other endogenous OCT substrates. The geometric mean plasma cimetidine concentration-time profile is shown in Figure S1. The geometric mean metformin plasma concentrations were consistently higher in the presence relative to the absence of cimetidine (Figure 1C). Metformin AUC_(0-24 h)_ and C_max_ were significantly increased by 24% (*p* = 0.019) and 36% (*p* = 0.004), respectively, in the presence of cimetidine (Figure 1D, E). A_e(0-24 h)_ showed a modest increase (18%) upon cimetidine exposure (Figure 1F), resulting in a small decrease in CL_r,a_ (8%, *p* = 0.005) (Figure 1G). Metformin AUC_(0-4 h)_ significantly increased by 28% in the presence of cimetidine, while CL_r,b_ was unaffected (Table 1). The profiles were comparable between females and males during both the baseline (Figure S2A) and cimetidine exposure (Figure S3A) arms; no sex-dependent differences were detected (Figures S2B-E and Figures S3B-E).

**Table 1.** Pharmacokinetics of OCT substrates during baseline and cimetidine exposure arms in healthy adults (n=16).

| OCT substrate | Pharmacokinetic Measure | Baseline | Cimetidine exposure | %Change (↑/↓)<br>(Cimetidine exposure/baseline) | p-value |
| --- | --- | --- | --- | --- | --- |
| Metformin | AUC<sub>(0-24 h)</sub><br>(μg*h/L) | 830.8<br>(710.6-971.3) | 1029.4<br>(875.0-1211.0) | 24% (↑) | 0.019* |
|  | C<sub>max</sub><br>(μg *h/L) | 123.7<br>(106.9-143.2) | 167.9<br>(144.9-194.6) | 36% (↑) | 0.004** |
|  | Ae<sub>(0-24 h)</sub><br>(μg) | 25622<br>(22769-28833) | 30274<br>(27843-32917) | 18% (↑) | 0.032* |
|  | CL<sub>r,a</sub><br>(L/h) | 30.6<br>(27.2-34.5) | 28.2<br>(25.3-31.4) | 8% (↓) | 0.005** |
|  | AUC<sub>(0-4 h)</sub><br>(μg*h/L) | 395.5<br>(341.6-458.0) | 507.2<br>(434.6-591.2) | 28% (↑) | 0.045* |
|  | Ae<sub>(0-4 h)</sub><br>(μg) | 9812<br>(7947-12114) | 10222<br>(7862-13291) | 4% (↑) | 0.658 |
|  | CL<sub>r,b</sub><br>(L/h) | 24.3<br>(18.9-31.3) | 20.1<br>(15.7-25.7) | 17% (↓) | 0.267 |
| N1-Methylnicotinamide | AUC<sub>(0-24 h)</sub><br>(μg*h/L) | 579.2<br>(509.8-658.1) | 530.6<br>(457.2-615.9) | 8% (↓) | 0.398 |
|  | C<sub>max</sub><br>(μg *h/L) | 50.5<br>(43.47-58.63) | 31.3<br>(26.7-36.7) | 38% (↓) | 0.004** |
|  | Ae<sub>(0-24 h)</sub><br>(μg) | 7439<br>(6516-8438) | 7415<br>(6516-8438) | 9% (↓) | 0.902 |
|  | CL<sub>r,a</sub><br>(L/h) | 12.6<br>(11-14.4) | 14<br>(12.4-15.7) | 1% (↑) | 0.298 |
|  | AUC<sub>(0-4 h)</sub><br>(μg*h/L) | 127.5<br>(110.1-147.6) | 74.9<br>(66.5-84.3) | 41% (↓) | <0.001*** |
|  | Ae<sub>(0-4 h)</sub><br>(μg) | 1642<br>(1368-1970) | 1170<br>(979-1397) | 29% (↓) | 0.041* |
|  | CL<sub>r,b</sub><br>(L/h) | 12.5<br>(10.4-15.1) | 15.3<br>(12.8-18.2) | 22% (↑) | 0.117 |
| N1-Methyladenosine | AUC<sub>(0-24 h)</sub><br>(μg*h/L) | 42.7<br>(34.9-52.1) | 40.7<br>(34.6-47.7) | 5% (↓) | 0.485 |
|  | C<sub>max</sub> | 2.6 | 2.6 | No change | 0.579 |
| | ($\mu\text{g} \cdot \text{h/L}$) | (2.1-3.2) | (2.2-3.0) | | |
| | **Ae<sub>(0-24 h)</sub>**<br>($\mu\text{g}$) | 174.9<br>(116.0-263.9) | 167.4<br>(105.6-265.1) | 4% (↓) | 0.943 |
|  | **CL<sub>r,a</sub>**<br>(L/h) | 4.1<br>(2.6-6.5) | 4.1<br>(2.5-6.7) | No change | 0.810 |
| | **AUC<sub>(0-4 h)</sub>**<br>($\mu\text{g} \cdot \text{h/L}$) | 7.0<br>(5.6-8.7) | 6.6<br>(5.7-7.7) | 5% (↓) | 0.367 |
| | **Ae<sub>(0-4 h)</sub>**<br>($\mu\text{g}$) | 21.9<br>(11.8-40.6) | 31.5<br>(16.8-59.1) | 44% (↑) | 0.669 |
|  | **CL<sub>r,b</sub>**<br>(L/h) | 3.1<br>(1.6-6.0) | 4.7<br>(2.5-8.7) | 53% (↑) | 0.810 |
| **Isobutyryl-L-carnitine** | **AUC<sub>(0-24 h)</sub>**<br>($\mu\text{g} \cdot \text{h/L}$) | 1015<br>(838-1230) | 876<br>(770-1238) | 4% (↓) | 0.806 |
| | **C<sub>max</sub>**<br>($\mu\text{g} \cdot \text{h/L}$) | 56.8<br>(46.3-69.7) | 53.7<br>(41.7-69.1) | 6% (↓) | 0.848 |
| | **AUC<sub>(0-4 h)</sub>**<br>($\mu\text{g} \cdot \text{h/L}$) | 151.9<br>(121.7-189.4) | 132<br>(108.6-160.3) | 16% (↓) | 0.076 |

AUC<sub>(0-24 h)</sub>, area under the plasma-concentration time curve from time zero to 24 h; AUC<sub>(0-4 h)</sub>, area under the plasma-concentration time curve from time zero to 4 h; C<sub>max</sub>, maximum plasma concentration from time zero to 24 h; Ae<sub>(0-24 h)</sub>, amount excreted into the urine from 0-24 h; Ae<sub>(0-4 h)</sub>, amount excreted into the urine from 0-4 h; CL<sub>r</sub>, renal clearance. CL<sub>r,a</sub> was calculated by dividing Ae<sub>(0-24 h)</sub> by AUC<sub>(0-24 h)</sub>. CL<sub>r,b</sub> was calculated by dividing Ae<sub>(0-4 h)</sub> by AUC<sub>(0-4 h)</sub>. The data are presented as a geometric mean (90% confidence Interval). Statistical analysis was conducted using two-tailed paired t-tests: *p-value* < 0.05 (\*) and < 0.01 (\*\*) compared to baseline.

### Effect of cimetidine on the pharmacokinetics of potential OCT substrates

The hypothesized interaction of NMN and NMA (purported OCT2 biomarkers) with cimetidine is expected to reduce CL_r_ of NMN and NMA due to OCT2 inhibition in renal proximal tubule cells (Figure 2A). NMN plasma concentrations were significantly lower in the cimetidine exposure arm compared to the baseline arm (Figure 2B). There was no change in AUC_(0-24 h)_ (Figure 2D), whereas NMN C_max_ decreased by 38% (*p* = 0.004) (Figure 2E). Cimetidine had no effect on A_e(0-24 h)_ and CL_r,a_ of NMN (Figures 2F, G). NMN AUC_(0-4 h)_ decreased by 41% after cimetidine coadministration, while CL_r,b_ was unchanged (Table 1). NMA plasma concentrations (Figure 2C), AUC, C_max_, A_e_, and CL_r_ were comparable between baseline and cimetidine exposure during both the 0-4 h and 0-24 h time intervals (Table 1, Figures 2H-K).

**Figure 2.**
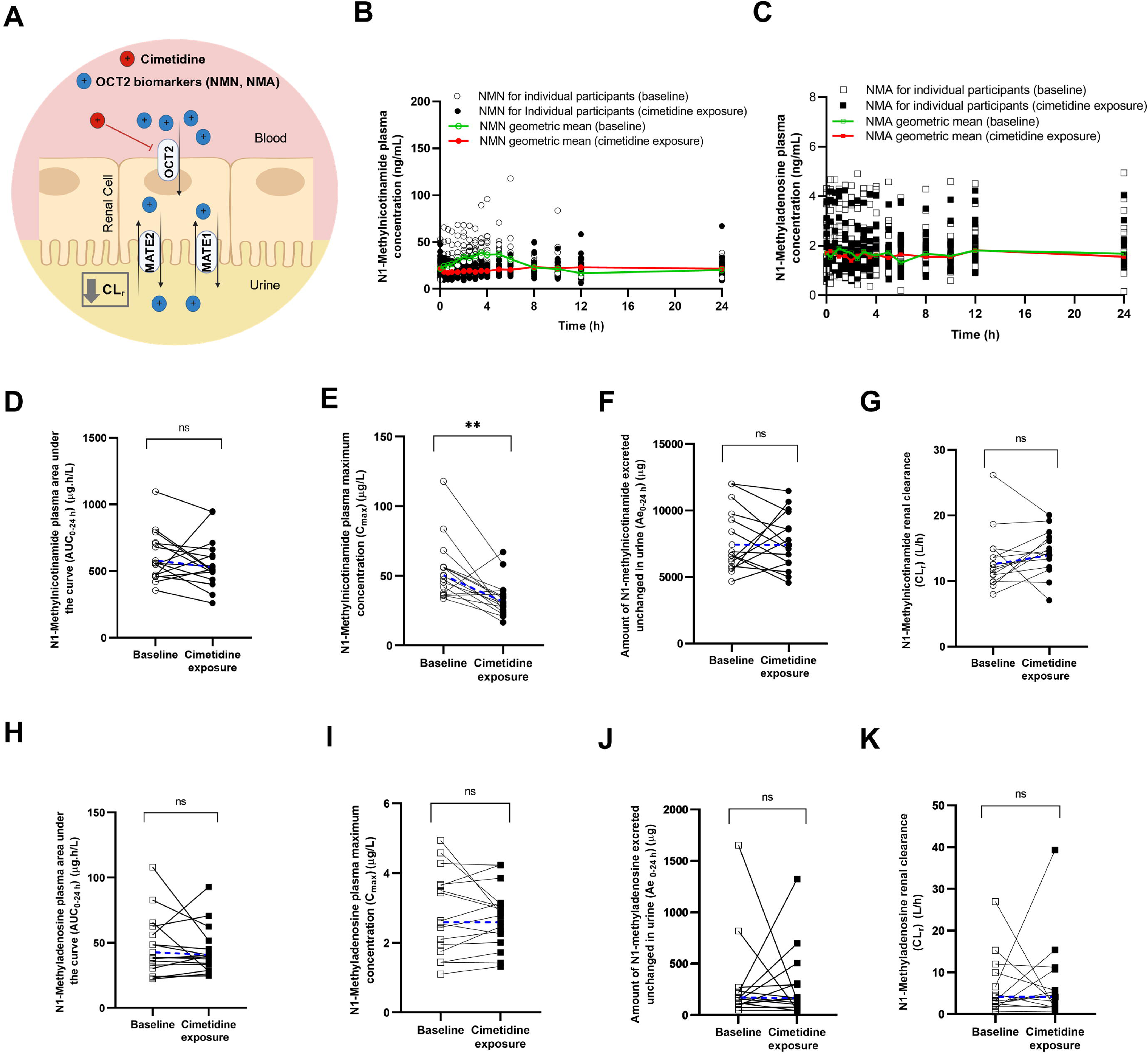
Effect of cimetidine co-administration on known endogenous OCT2 biomarkers, N1-methylnicotinamide (NMN) and N1-methyladenosine (NMA). Schematic representation of transport processes for the potential OCT2 biomarkers (NMN and NMA) and the hypothesized interaction with cimetidine in renal proximal tubule cells (A). Plasma concentration-time profiles of NMN (B) and NMA (C) in the metformin alone (baseline, open symbol) and metformin plus cimetidine (cimetidine exposure, solid symbol) arms among healthy adult participants (n=16). The plasma area under the curve from 0-24 h (AUC_0-24 h_) (D and H), plasma maximum concentration (C_max_) (E and I), amount excreted into the urine from 0-24 h (Ae_0-24 h_) (F and J), and renal clearance (CL_r_) (G and K) of NMN and NMA, respectively, in the metformin alone (baseline, open symbols) and metformin plus cimetidine (cimetidine exposure, solid symbols) arms among healthy adult participants (n=16). One participant was excluded from the Ae_0-24 h_ and CL_r_ analysis due to difficulty voiding urine. In Figure 2G and K, CL_r_ was calculated based on (0-24 h) time interval. Symbols connected by solid black lines represent individual values, while symbols connected by dashed blue lines represent geometric means. Statistical analysis was conducted using two-tailed paired t-tests: *p-value* > 0.05 (ns, not significant) and < 0.01 (**). Figure 2A was created using BioRender.

The hypothesized interaction of IBC (purported OCT1 biomarker) with cimetidine is expected to reduce IBC plasma systemic exposure due to OCT1 inhibition in hepatocytes (Figure 3A). IBC plasma concentrations decreased during the first 4 hours following cimetidine administration but did not reach statistical significance (Figure 3B). AUC_(0-24 h)_, AUC_(0-4 h)_, and C_max_ did not differ between the two arms of the study (Table 1, Figure 3C, D). Using untargeted analysis of plasma samples described in the Supplementary File, we detected a peak for IBC (*m/z* 233.1540, retention time 9.5 minutes), which matched IBC targeted analysis data; there was no significant alteration in IBC plasma concentrations between the two study arms. Two additional features with the same high-resolution *m/z* value (233.1542) as that of IBC were detected but had different retention times (22.3 and 23.5 minutes). Both features showed a significant reduction in plasma concentrations by 36% (*p* = 0.042) and 25% (*p* = 0.022), respectively, upon cimetidine treatment (Figures S4A, B). Targeted analysis of glycine betaine, tryptophan, N1-methyl nicotinic acid, choline, acetylcholine, and carnitine as potential OCT substrates showed no significant differences in relative plasma concentrations between study arms (Figures S5A-F).

**Figure 3.**
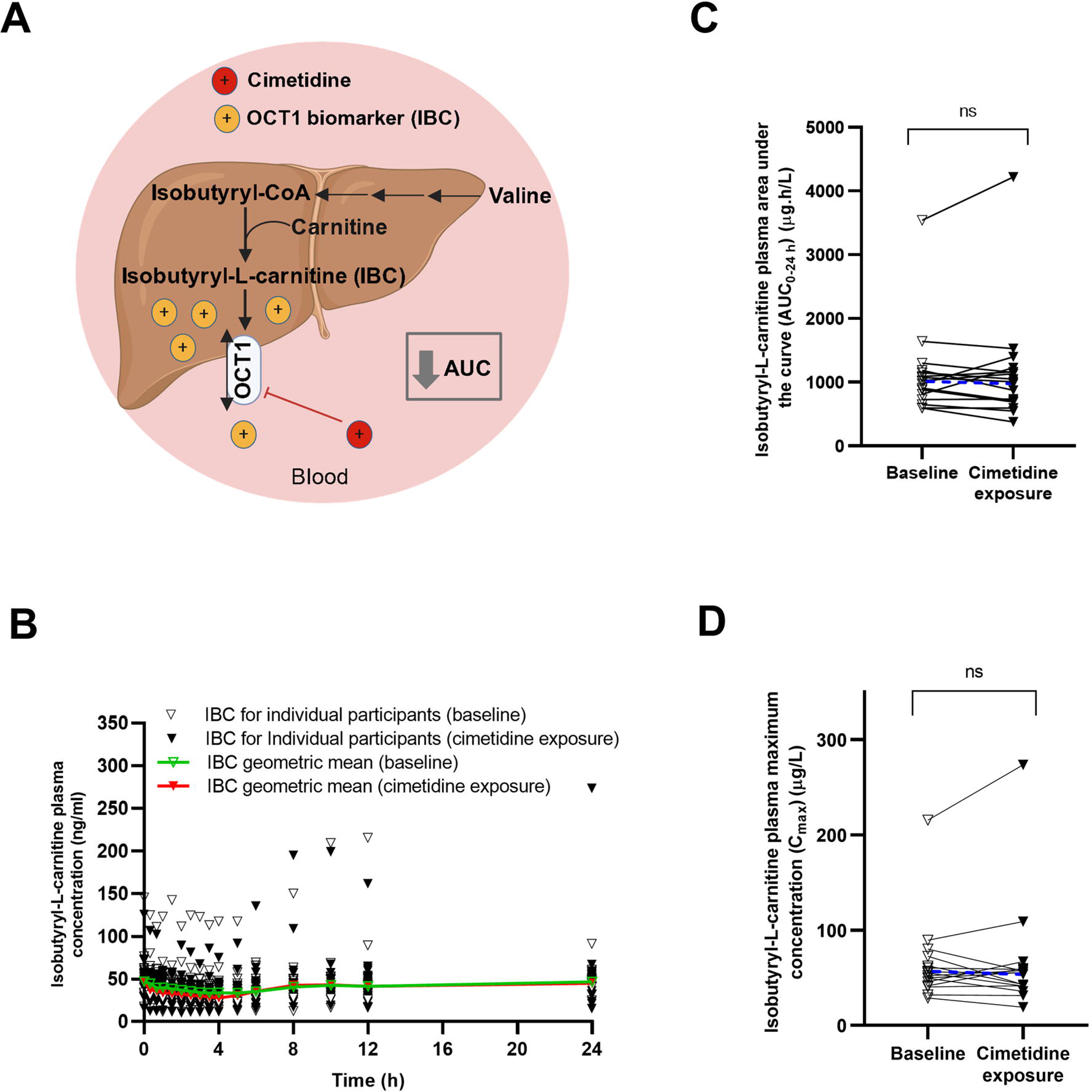
Effect of cimetidine co-administration on known endogenous OCT1 biomarker, isobutyryl-L-carnitine (IBC). Schematic representation of transport processes for the potential OCT1 biomarker (IBC) and the hypothesized interaction with cimetidine in hepatocytes (A). Plasma concentration-time profiles (B), plasma area under the curve from 0-24 h (AUC_0-24 h_) (C), and plasma maximum concentration (C_max_) (D) of IBC in the metformin alone (baseline, open symbols) and metformin plus cimetidine (cimetidine exposure, solid symbols) arms among healthy adult participants (n=16). Symbols connected by solid black lines represent individual values, while symbols connected by dashed blue lines represent geometric means. Statistical analysis was conducted using two-tailed paired t-tests: *p-value* > 0.05 (ns, not significant). Figure 3A was created using BioRender.

### Predicted contributions of hepatic OCT1 and renal OCT2 to the pharmacokinetic cimetidine-metformin interaction

The PBPK model successfully captured the cimetidine-metformin interaction, as the observed metformin concentrations were within 5^th^ and 95^th^ percentiles of simulated concentrations for both study arms (Figure 4A). The model was mechanistic and captured the various pharmacokinetic endpoints within 2-fold of the observed values (Table 2). The contribution of hepatic OCT1 was predicted to be 21% when renal OCT2 was omitted from the metformin model (Figure 4B), whereas the contribution of renal OCT2 was 10% when hepatic OCT1 was omitted (Figure 4C). The model predicted a reduction in metformin hepatic concentration by 27% following cimetidine administration (Figure 4D). The cimetidine PBPK model successfully captured the observed data (Figure S6A). The metformin PBPK model successfully captured metformin plasma concentrations at the subtherapeutic dose (50 mg) reported by Nguyen et al. (Figure S6B),^16^ as well as those for a range of metformin doses (10-500 mg) administered with and without cimetidine (Figures S6D-F).^9,17,18^ PBPK model predicted a reduction in metformin hepatic concentration by increasing biliary clearance (CL_bile_) (Figures S7A, B) regardless of the presence of cimetidine, without altering metformin plasma concentrations (Figure S7C).

**Figure 4.**
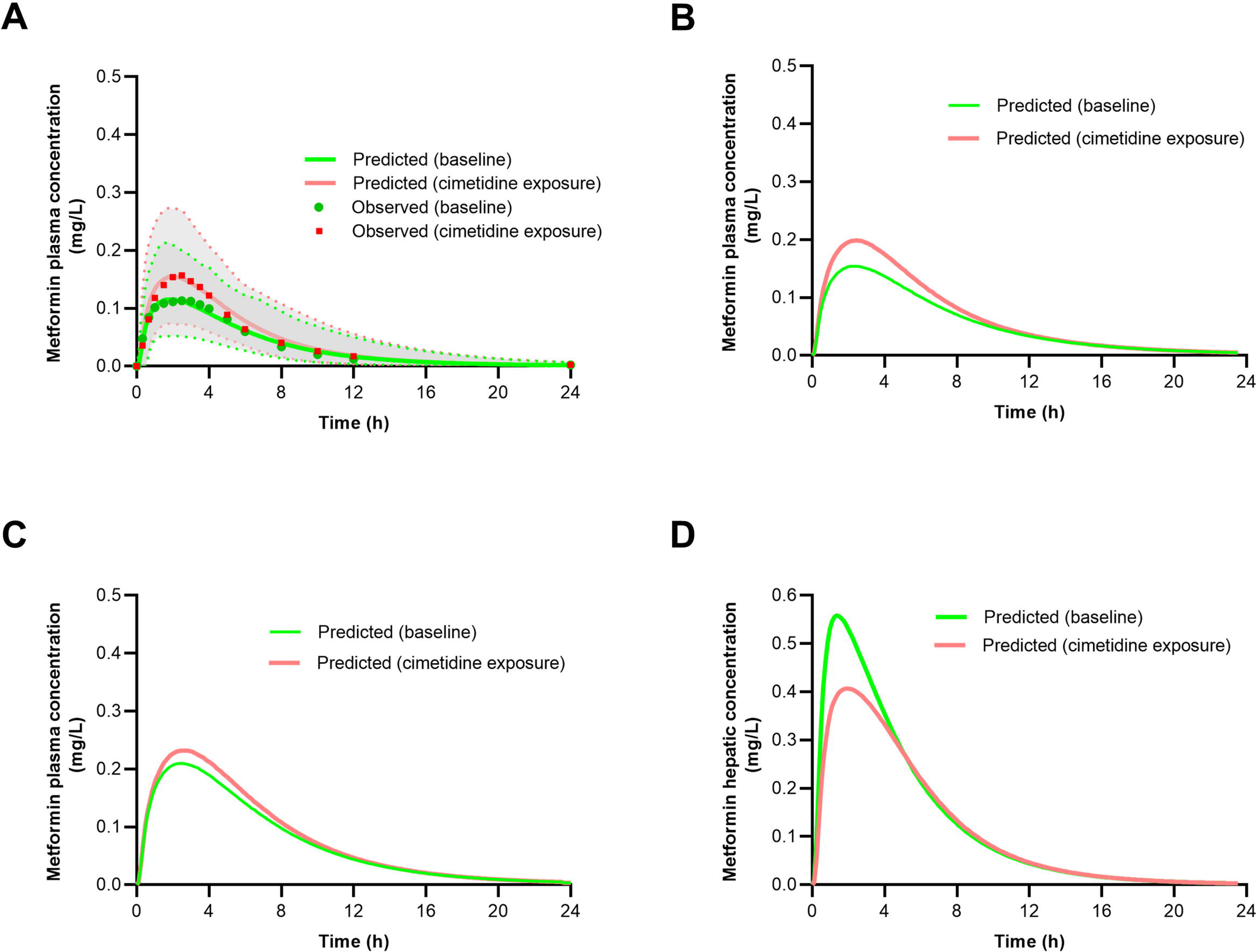
Physiologically based pharmacokinetic (PBPK) model-predicted contribution of hepatic OCT1 versus renal OCT2 inhibition to the cimetidine-metformin interaction. The PBPK model accurately predicted plasma concentration-time profiles of metformin without (baseline) and with cimetidine (cimetidine exposure) (A). The model predicted pharmacokinetic profiles indicating individual contributions of hepatic OCT1 (B) and renal OCT2 (C) to the pharmacokinetic cimetidine-metformin interaction. Arithmetic means of predicted metformin plasma concentrations are depicted as solid green (baseline) and solid red (cimetidine exposure) lines. The model-predicted hepatic concentration profile of metformin alone (baseline) and with cimetidine (cimetidine exposure) (D). The 5^th^ and 95^th^ percentiles of the predicted metformin plasma concentrations are illustrated with the shaded areas constrained by green (baseline) and red (cimetidine exposure) dashed lines. The observed data are represented as green dots (baseline) and red squares (cimetidine exposure).

**Table 2.** Comparison of pharmacokinetic measures and contribution of hepatic OCT1 and renal OCT2 in the presence and absence of cimetidine between observed (current study) and simulated (PBPK model) data.

| Pharmacokinetic measure | % Change (↑/↓)<br>(cimetidine exposure/baseline)<br>Observed data | % Change (↑/↓)<br>(cimetidine exposure/baseline)<br>Simulated data | Ratio<br>(simulated/observed) |
| --- | --- | --- | --- |
| **AUC<sub>(0-24 h)</sub>** | 24% (↑) | 26% (↑) | 1.1 |
| **C<sub>max</sub>** | 36% (↑) | 34% (↑) | 0.94 |
| **A<sub>e(0-24 h)</sub>** | 18% (↑) | 15% (↑) | 0.83 |
| **CL<sub>r</sub>** | 8% (↓) | 10% (↓) | 1.25 |
| **Hepatic concentration** | N/A | 27% (↓) | --- |
| **Hepatic OCT1 alone**<br>**(AUC<sub>(0-24 h)</sub>)** | 16% (↑) | 21% (↑) | 1.31 |
| **Renal OCT2 alone**<br>**(AUC<sub>(0-24 h)</sub>)** | 8% (↑) | 10% (↑) | 1.25 |

AUC<sub>(0-24 h)</sub>, area under the plasma-concentration time curve from time zero to 24 h; C<sub>max</sub>, maximum plasma concentration from time zero to 24 h; A<sub>e(0-24 h)</sub>, amount excreted into the urine from 0-24 h; CL<sub>r</sub>, renal clearance; N/A, not applicable. CL<sub>r</sub> was calculated based on (0-24 h) time interval in both observed and predicted data.

### Effects of *OCT* and *MATE* genetic variants on metformin pharmacokinetics and interaction with cimetidine

Nine *OCT1*, three *OCT2*, four *MATE1*, and four *MATE2K* genetic variants were selected for analysis as these were found in at least three study participants. Individuals were assigned to reference and alternative (individual or combined homozygous and heterozygous) genotype groups as shown in Table S7. Out of these genetic variants, one *OCT1*, two *OCT2*, and two *MATE2K* variants resulted in statistical significance of cimetidine-metformin interaction and magnitude (% change after cimetidine treatment) in pharmacokinetic measures of metformin assigned to each variant as detailed in Figure 5. Individuals homozygous for the *OCT1* c.480G>C (p.Leu160Phe, rs683369) variant showed a statistically significant increase in metformin AUC_(0-24 h)_ and C_max_ of 33% (*p* = 0.025) and 48% (*p* = 0.006), respectively, with cimetidine treatment compared to no significant change in pharmacokinetic measures compared to heterozygous individuals (Figures 5A, B). Participants homozygous for *OCT2* c.518+32G>C (rs2774230) or c.1506C>T (p.Val502=, rs316003) demonstrated a statistically significant increase in AUC_(0-24 h)_ by 35% (*p* = 0.013) and 27% (*p* = 0.022), respectively, and reduction by 9% in metformin CL_r,a_ when co-administered with cimetidine compared to a moderate change in AUC_(0-24 h)_ and CL_r,a_ for individuals who are heterozygous for these variants (Figures 5C, D). *MATE2K* genetic variants c.-130C>T (rs12943590) and c.345C>T (rs4924792) resulted in 33% increase and 9% decrease in metformin AUC_(0-24 h)_ and CL_r,a_, respectively, in the cimetidine exposure arm for the reference genotype groups with no statistically significant change in pharmacokinetic measures when compared with the genotype group comprising individuals with one or two variant alleles (Figures 5G, H). While *MATE1* c.-66T>C (rs2252281) did not affect the cimetidine-metformin interaction, this variant was associated with higher metformin AUC_(0-24 h)_ in the homozygous variant group TT by 1.42-fold (*p* = 0.042) compared to the heterozygous group TC in the baseline arm (Figures 5E, F).

**Figure 5.**
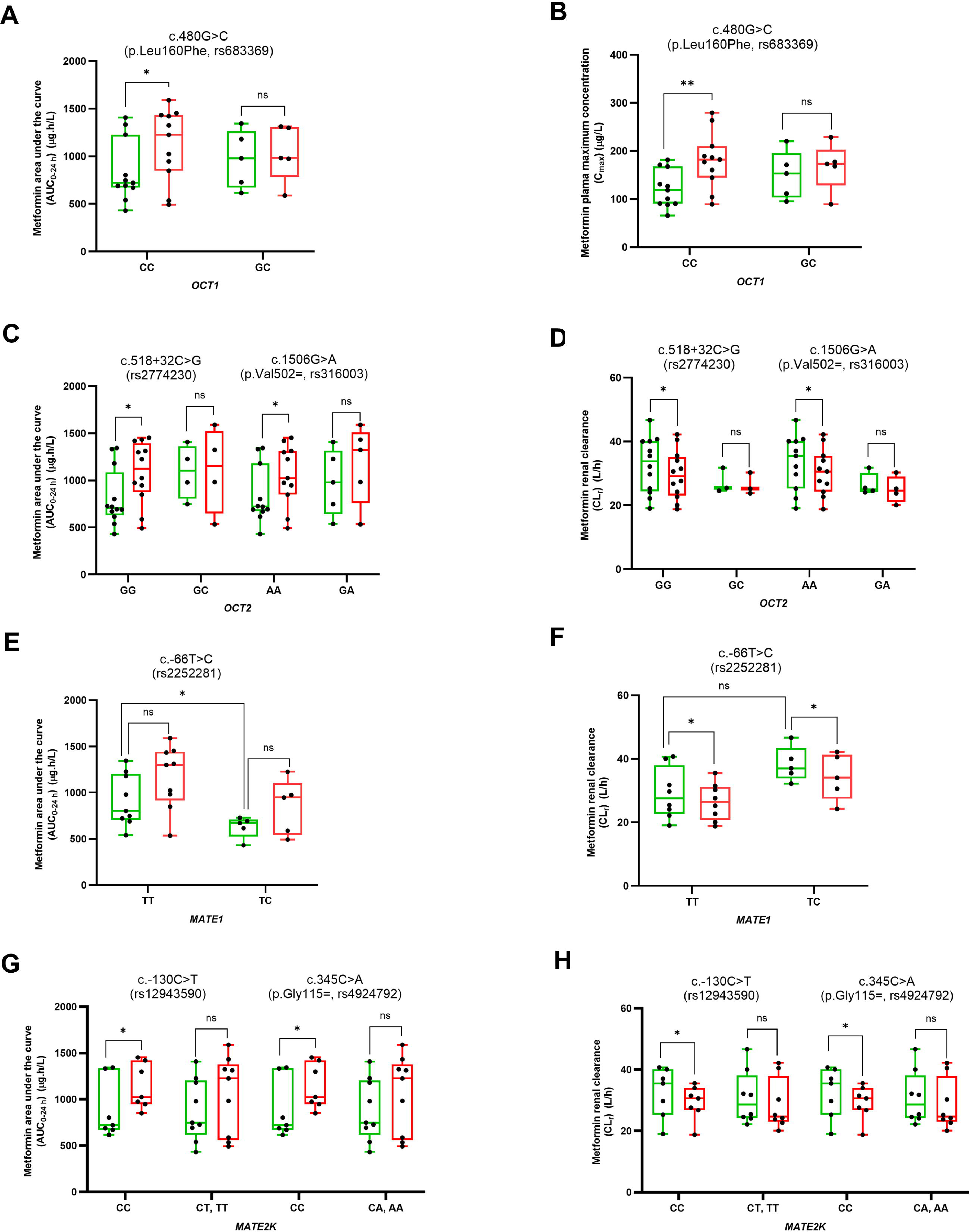
Effect of *OCT1*, *OCT2*, *MAET1*, and *MATE2K* genetic variants on the cimetidine-metformin interaction. The effect of *OCT1 genetic variants on* plasma area under the curve from 0-24 h (AUC_0-24 h_) (A) and plasma maximum concentration (C_max_) (B), *OCT2* genetic variants on plasma area under the curve from 0-24 h (AUC_0-24 h_) (C) and renal clearance (CL_r_) (D), *MATE1* genetic variants *on* plasma area under the curve from 0-24 h (AUC_0-24 h_) (E) and renal clearance (CL_r_) (F) and *MATE2K* genetic variants on plasma area under the curve from 0-24 h (AUC_0-24 h_) (G) and renal clearance (CL_r_) (H) of metformin when administered alone (baseline, green) and in combination with cimetidine (cimetidine exposure, red) in healthy adults (n=16). One participant was excluded from the CL_r_ analysis due to difficulty voiding urine. In Figures 5D, F, and H, CL_r_ was calculated based on (0-24 h) time interval. Statistical analysis was conducted for each group carrying a similar genotype with and without cimetidine using two-tailed paired t-tests: *p-value* > 0.05 (ns, not significant), < 0.05 (*) and < 0.01 (**). Statistical analysis was conducted for metformin disposition in baseline arm according to genotype using unpaired t-tests followed by the Mann-Whitney test: *p-value* > 0.05 (ns, not significant) and < 0.05 (*).

*OCT1* (c.411+232C>G (rs4709400), c.412-43T>G (rs4646272), c.516-26C>T (rs45584532), c.516-26C>T (rs45584532), c.1498+43C>T (rs2297374), c.1222A>G (p.Met408Val, rs628031), c.1276+9_1276+16delTGGTAAGT (rs2114790299), and c.1260_1262delATG (p.Met420del, rs72552763)), *OCT2* (c.390G>T (p.Thr130=, rs624249)), *MATE1* (c.499-4G>A (rs2247436), c.499-12G>C (rs2247437), c.922-158G>A (rs2289669)), and *MATE2K* (c.885C>T (p.Tyr331=, rs4925042) and c.841+14G>C (rs12942065)) genetic variants were not associated with metformin pharmacokinetics or cimetidine-metformin interaction (Table S7).

## DISCUSSION

OCTs are crucial in mediating the absorption, distribution, and clearance of metformin and can influence metformin pharmacokinetics and potentially, glucose-lowering effects.^19,20^ OCT2 is mainly associated with metformin pharmacokinetics, whereas OCT1 plays a key role in metformin response.^4^ Shu et al. further demonstrated individuals carrying a heterozygous or homozygous reduced-function *OCT1* allele exhibited higher metformin systemic plasma exposure compared to those homozygous for the reference *OCT1* allele.^21^ OCT-mediated DDIs should be considered when co-administering an OCT inhibitor such as cimetidine with metformin. While multiple studies only focused on the effect of pharmacokinetic interactions on metformin CL_r_ mediated by OCT2 inhibition, the effect on hepatic OCT1 remains underappreciated.

We conducted a pharmacokinetic cimetidine-metformin interaction study where a subtherapeutic dose of metformin was selected to minimize toxicity risk and reduce inter-individual variability related to nonlinear absorption kinetics associated with therapeutic doses.^22,23^ Our data suggested that the cimetidine-mediated increase in metformin exposure (24%) accounted for the interactions with both hepatic OCT1 and renal OCT2, while the decrease in CL_r_ (8%) accounted for the interaction with renal OCT2, further suggesting that hepatic OCT1 accounted for the remaining 16% of the interaction. Previously reported cimetidine-metformin DDIs involving healthy adults showed metformin CL_r_ decreased by 27-45% when oral metformin (250-500 mg) was administered with cimetidine.^9,17,18^ This moderate effect on metformin CL_r_ could be due to multiple reasons. First, cimetidine can inhibit other cation transporters, including OCT1 and MATE1/2K.^11,24^ Second, the single cimetidine dose (400 mg) used in our study differed from the cimetidine regimen used in other studies. Somogyi et al. administered cimetidine (400 mg) twice daily for 6 days before administering metformin^9^, while Wiebe et al. administered 400 mg of cimetidine before metformin administration, with additional cimetidine doses administered 4, 8, 12, 24, and 36 h later.^17^ Cimetidine plasma concentrations in our study were at least 4 times lower than the reported IC_50_ for OCT2, which range from 23.6-207 µM.^11,24,25^ A recent study by Koishikawa et al. used a similar cimetidine dosing regimen reported a 15% reduction in metformin CL_r_ though this reduction was not statistically significant suggesting weak renal interaction at 400 mg cimetidine dose consistent with our findings.^26^ Third, equal numbers of males and females were enrolled in our study; whereas only healthy males participated in previous cimetidine-metformin clinical studies.^9,17,18^ However, this is less likely to be the reason considering no sex difference was observed in our study. Finally, genetic variation in the transporter genes responsible for metformin disposition may affect the outcome of the cimetidine-metformin DDI due to altered transporter activity and the subsequent effect on fraction transported (ft) by individual protein. Inhibition of metformin renal tubular secretion by cimetidine appears to be dependent on genetic variation in the *OCT2* gene. Wang et al. showed that metformin CL_r_ was significantly reduced “TT” carriers of 808G>T (p.Ser270Ala, rs316019) variant compared to “GG” carriers,^18^ thus “TT” carriers were less sensitive to cimetidine inhibition which is consistent with our observations with other *OCT2* genetic variants (rs2774230 and rs316003).^18^ Notably, the 808G>T (p.Ser270Ala, rs316019) variant was detected in only two participants, which limited the ability to perform statistical analysis. Although the effect of *OCT1* c.480G>C (p.Leu160Phe, rs683369) on metformin pharmacokinetics has not been reported, this variant was associated with metformin response in patients with polycystic ovarian syndrome, where heterozygous “GC” individuals had higher oral glucose tolerance compared to carriers of the homozygous variant allele.^27^ Our data did not reveal any associations of *MATE1* and *MATE2K* genetic variants with metformin pharmacokinetics at baseline which is in agreement with reports showing no association with an intronic *MATE1* variant (rs2289669) or *MATE2K* (rs12943590).^19^

To further evaluate the contribution of renal OCT2 and hepatic OCT1 to the cimetidine-metformin DDI, we conducted a full pharmacokinetic analysis of potential biomarkers of OCT2 (NMN and NMA) and OCT1 (IBC). NMN is mainly synthesized in the liver from nicotinamide,^28^ whereas NMA is post-transcriptional modification of RNA though methylation of adenosine moiety.^29^ Both the metabolites are substrates of OCT2 and are eliminated in urine, with a CL_r_ 2-6 fold higher than GFR, suggesting the involvement of renal secretion.^30,31^ Administration of OCT inhibitors such as cimetidine, trimethoprim, and pyrimethamine resulted in a reduction in CL_r_ of NMN^13,30,32^ and NMA.^14^ However, we did not observe a reduction in NMN and NMA CL_r_ in the cimetidine exposure arm relative to baseline, which further suggests a minimum interaction in the kidney with a single 400 mg dose of cimetidine in alignment with Koishikawa et al. recent data reported.^26^ We also observed a decrease in plasma NMN concentration upon cimetidine administration, consistent with that observed using different OCT inhibitors.^13,30,32^ NMN is a hepatic OCT1 substrate, with an 11-fold increase in NMN uptake by OCT1-transfected HEK293 cells compared to mock HEK293 cells.^33^ Cimetidine, acting as an OCT1 inhibitor, may impede the release of NMN into the plasma, leading to NMN accumulation in the liver. In our study, NMN captured an interaction only in the liver from 0 to 4 h (reduced NMN plasma concentration without alteration in CL_r_), suggesting NMN as a potential biomarker for hepatic OCT1. IBC, a potential OCT1 biomarker, is formed in hepatocytes as a metabolite of valine and isobutyryl-CoA.^34^ In our study, IBC did not predict an interaction in the liver because no significant change was observed in the pharmacokinetic measures between the two arms of the study at both the 0-4 h and 0-24 h time intervals, suggesting that IBC may not be a reliable OCT1 biomarker for weak interactions. However, untargeted analysis revealed two other features matching IBC *m/z* value that significantly decreased after cimetidine exposure. Those features may be related to IBC analogues and may have a potential role as OCT1 biomarkers but require further structural characterization.

Multiple PBPK models have been developed to gain mechanistic insight into the cimetidine-metformin interaction.^35^ However, these models did not consider the possible contribution by hepatic OCT1. We established and verified a model using our data and data from other studies. Our model successfully predicted metformin plasma concentrations at a subtherapeutic dose (50 mg).^16,17^ Model performance remained robust at therapeutic doses (250 and 500 mg),^917,18^ regardless of the presence of cimetidine. Accordingly, we have confidence in the reliability of our model for assessing the contributions of hepatic OCT1 and renal OCT2 to the cimetidine-metformin interaction. Further investigation is needed to assess the impact of cimetidine on metformin pharmacodynamics and efficacy because our PBPK model predicted a larger contribution by hepatic OCT1 (21%) compared to renal OCT2 (10%) and a 27% reduction in metformin hepatic concentration following cimetidine administration. CL_bile_ sensitivity analysis showed primarily effects on metformin hepatic concentrations, suggesting a potential impact on metformin response. Considering that CL_bile_ of metformin can be mediated through MATE1 expressed at the bile canalicular membrane of hepatocytes, Stocker et al. showed that *MATE1* variants resulted in higher metformin response in patients with diabetes.^36^

We would like to acknowledge key limitations of this study. First, calibration curve standards and QC samples were prepared using a matrix surrogate or neat solvent, given that we are monitoring endogenous compounds. However, to account for matrix effects, wherever possible, we compared the metabolite of interest with its respective stable isotope labeled (internal standard). Additionally, the study was not powered to detect sex differences. Second, only a small number of participants were assigned to genotype groups for analysis of genetic variants of different transporters, which may have limited our ability to detect significant effects on metformin pharmacokinetics. Furthermore, some SNPs may have been found, but not in enough participants, other SNPs may not have been detected at all, and WES does not detect some genetic variants of potential interest which are located deeper in introns such as *OCT1* c.1386-2964C>A (rs622342). Lastly, although the pharmacogenetic portion of this investigation had limitations, our findings corroborate that the ft by individual transporters can be affected by genetic variation in one or more of these transporters, leading to varying magnitudes of DDI between cimetidine and metformin. Prospective clinical studies are needed to confirm these findings, assess additional variants, and assess the impact of variants in diverse populations using different OCT inhibitors.

In summary, our investigation of the cimetidine-metformin DDI provided comprehensive insights into the transporters involved. We showed for the first time a potential role for hepatic OCT1, in addition to renal OCT2, in this DDI. Because most OCT inhibitors lack selectivity, the contribution by OCT1 to the hepatic disposition of metformin should not be underestimated given that the liver is the primary site of action. Our PBPK model successfully captured the respective contribution of both hepatic OCT1 and renal OCT2. The use of endogenous OCT2 metabolites allowed for the assessment of renal DDIs, indicating minimal renal OCT2 involvement in the cimetidine-metformin interaction, particularly when a subtherapeutic metformin dose was administered with a single dose of cimetidine. Further research is needed to evaluate the potential effect of cimetidine on metformin efficacy.

## STUDY HIGHLIGHTS

1. **What is the current knowledge on the topic?** The pharmacokinetic interaction between cimetidine (precipitant) and metformin (object) is primarily mediated through organic cation transporters (OCTs) expressed in liver (OCT1) and kidney (OCT2). Metformin pharmacokinetics are driven by hepatic OCT1 and renal OCT2. However, the contributions of OCT1 and OCT2 to the interaction, as well as the impact of genetic variation on the interaction and metformin pharmacokinetics, remain unknown.
2. **What question did this study address?** We investigated the contributions of hepatic OCT1 and renal OCT2 to the cimetidine-metformin interaction. Next, we examined the utility of physiologically based pharmacokinetic (PBPK) modeling and potential biomarkers of OCT1 (isobutyryl-L-carnitine) and OCT2 (N1-methylnicotinamide and N1-methyladenosine) in detecting the interaction between cimetidine and metformin. Finally, we characterized the effect of variants in the *SLC22A1* and *SLC22A2* genes encoding OCT1 and OCT2 variants on metformin pharmacokinetics and the interaction with cimetidine.
3. **What does this study add to our knowledge?** Cimetidine inhibits both hepatic OCT1 and renal OCT2, affecting metformin pharmacokinetics and potentially altering metformin efficacy. N1-methylnicotinamide, N1-methyladenosine, and isobutyryl-L-carnitine may not serve as reliable OCT biomarkers for weak interactions. However, N1-methylnicotinamide shows promising potential as a biomarker for OCT1. Genetic variants in *OCT* and *MATE* can confound the magnitude of the cimetidine-metformin interaction.
4. **How might this change clinical pharmacology or translational science?** Results are essential for evaluating the sensitivity of some proposed biomarkers for OCT1 and OCT2 in predicting drug-drug interactions. This study highlighted a potential reduction in metformin efficacy when co-administered with cimetidine.

## ACKNOWLEDGEMENTS

The authors thank Ms. Deena Hadi, Ms. Maxey Cherel, and Mr. Ryan Davy for their expert assistance in the completion of the clinical study. The authors would also like to thank Dr. J. Steven Leeder and Dr. Kelsee Halpin for their input on genotyping assay selection.

## AUTHOR CONTRIBUTIONS

Participated in research design: A.S.A., M.F.P, E.C.B, A.G, and B.P.

Conducted experiments: A.S.A. and D.K.S.

Performed data analysis: A.S.A., D.K.S., E.C.B, and B.P.

Wrote or contributed to the writing of the manuscript: A.S.A., D.K.S., A.T., M.F.P, E.C.B, A.G, and B.P.

## SUPPLEMENTARY MATERIALS

### METHODS

**Whole exome sequencing**

**Bioanalytical method for OCT substrate analysis in plasma and urine**

**Untargeted metabolomics analysis of plasma samples**

**Statistical analysis**

**PBPK model development and verification of the cimetidine-metformin interaction**

### RESULTS

**Bioanalytical method validation**

### SUPPLEMENTARY TABLES

**Table S1.** Inclusion and exclusion criteria for the clinical pharmacokinetic cimetidine-metformin interaction study.

**Table S2.** Demographic characteristics of the healthy adults who participated in the clinical pharmacokinetic cimetidine-metformin interaction study.

**Table S3.** Liquid chromatography (LC) methods and parameters for metformin, cimetidine, and potential OCT biomarkers.

**Table S4.** MS transitions, cone voltage, and collision energies of analytes of interest and internal standards used for multiple reaction monitoring (MRM)-based targeted assays.

**Table S5**. Metformin-specific parameters used for metformin PBPK model development.

**Table S6**. Cimetidine-specific parameters used in the metformin PBPK model.

**Table S7.** Effects of *OCT* and *MATE* genetic variants on the clinical pharmacokinetic cimetidine-metformin interaction.

### SUPPLEMENTARY FIGURE LEGENDS

**Figure S1.** Geometric mean plasma concentration-time profile of cimetidine after a single oral dose of 400 mg in healthy subjects (n=16). Symbols and error bars represent observed geometric means and 90% confidence intervals, respectively.

**Figure S2.** Association of sex on the mean plasma concentration-time profile (A), plasma area under the curve from 0-24 h (AUC_0-24 h_) (B), plasma maximum concentration (C_max_) (C), amount excreted into the urine from 0-24 h (Ae_0-24 h_) (D), and renal clearance (CL_r_) (E) of metformin between healthy female (pink) and male (blue) (n=16, 8 females, 8 males) in metformin alone arm (baseline). One male was excluded from the Ae_0-24 h_ and CL_r_ analysis due to difficulty voiding urine. In Figure S2E, CL_r_ was calculated based on (0-24 h) time interval. Symbols and error bars denote observed means and 90% confidence intervals, respectively. Statistical analysis was conducted using two-tailed paired t-tests: *p-value* > 0.05 (ns, not significant).

**Figure S3**. Association of sex with mean plasma concentration-time profile (A), plasma area under the curve from 0-24 h (AUC_0-24 h_) (B), plasma maximum concentration (C_max_) (C), amount excreted into the urine from 0-24 h (Ae_0-24 h_) (D), and renal clearance (CL_r_) (E) of metformin between healthy female (pink) and male (blue) (n=16, 8 females, 8 males) in metformin plus cimetidine arm (cimetidine exposure). One male was excluded from the Ae_0-24 h_ and CL_r_ analysis due to difficulty voiding urine. In Figure S3E, CL_r_ was calculated based on (0-24 h) time interval. Symbols and error bars denote observed means and 90% confidence intervals, respectively. Statistical analysis was conducted using two-tailed paired t-tests: *p-value* > 0.05 (ns, not significant).

**Figure S4.** Effect of cimetidine co-administration on the known potential endogenous OCT1 biomarker isobutyryl-L-carnitine (IBC) analogues. LC-MS peak areas of IBC analogues; *m/z* 232.1541 with retention time 22.3 (A) and *m/z* 232.1542 with retention time 23.5 (B) in pooled plasma (n=12) at 1.5, 2, 2.5, 3, and 3.5 h in the metformin alone (baseline, open symbols) and metformin plus cimetidine (cimetidine exposure, solid symbols) arms among healthy adults. Solid lines indicate individual values and dashed line represents geometric mean. Statistical analysis was conducted using two-tailed paired t-tests: *p-value* < 0.05 (*).

**Figure S5.** Effect of cimetidine co-administration on the potential OCT substrates. LC-MS/MS area ratio of glycine betaine (A), tryptophan (B), N1-methyl nicotinic acid (C), choline (D), acetylcholine (E), and carnitine (F) in the metformin alone (baseline, open symbols) and metformin plus cimetidine (cimetidine exposure, solid symbols) arms among healthy adults (n=16).

**Figure S6.** Physiologically based pharmacokinetic (PBPK) model predicted mean cimetidine plasma concentration-time profiles from our study (A) after 400 mg single dose of cimetidine. Metformin PBPK model for metformin subtherapeutic dose (50 mg) was reproduced from Nguyen et al. 2021 (B). Cimetidine-metformin PBPK model predicted mean metformin plasma concentration for metformin at 10 (C), 250 (D), and 500 mg (E, F) doses. Population prediction arithmetic means of metformin plasma concentration are shown as solid green (baseline) and solid red (cimetidine exposure) lines. 5^th^ and 95^th^ percentiles of the predicted metformin plasma concentrations are illustrated with the area constrained by green (baseline) and red (cimetidine exposure) dash lines. The observed data are shown as green dots (baseline) and red squares (cimetidine exposure).

**Figure S7.** Local sensitivity analysis of biliary secretion (CL_bile_) in metformin physiologically based pharmacokinetic (PBPK) model. The PBPK model predicted impact of CL_bile_ on metformin hepatic concentration when administered alone (baseline) (A) and in combination with cimetidine (cimetidine exposure) (B), and metformin plasma concentration (C).

