## Supplementary File for "Quantitative contributions of hepatic and renal organic cation transporters to the clinical pharmacokinetic cimetidine-metformin interaction"

**Affiliation:**

**METHODS**

**Whole exome sequencing**

Whole blood collected was placed into two vacutainer tubes containing ethylenediaminetetraacetic acid (~3 mL per tube). Blood in the first tube was divided into two 2-mL cryovials. The second tube was centrifuged at 4˚C at 2,000*g* for 10 minutes to separate the plasma. Upon collecting the plasma, a 2-mm plasma layer remained above the buffy coat. The residual volume (plasma layer, buffy coat, and red blood cells) was mixed gently several times, and 300 µL was transferred to a microcentrifuge tube. Tubes containing the whole blood and residual mixture were placed on ice and shipped to the Pharmacogenetics Core Laboratory at Children’s Mercy Hospital (Kansas City, MO) for DNA isolation. Whole exome sequencing (WES) was performed by the Children’s Mercy Genomics Core Service Center.

WES libraries were prepared according to the manufacturer’s standard protocols for Illumina TruSeq library preparation and IDT XGen Exome Enrichment. Cleaned, adapter-blocked pools were loaded on a NovaSeq6000 or NovaSeq X Plus with a run configuration of 151x8x8x151. Samples were sequenced to an average read depth of 15Gb for and average 100x read coverage. Read alignments and variant calling were conducted using the Dragen Bio-IT platform (v 3.10.4, Illumina). Variants were called with QUAL scores of ≥ 20. Variants of interest were captured using GATK v4.5.0.0^1^ and BCFTools v1.19^2^ with the genomic coordinates of the MANE transcripts for the genes of interest: *SLC22A1* [*OCT1*] (NM_003057.3), *SLC22A2* [*OCT2*] (NM_003058.4), *SLC47A1* [*MATE1*] (NM_018242.3), and *SLC47A2* [*MATE2K*] (NM_001099646.3). The MANE transcripts were also used as a feature filter for Variant Effect Predictor (VEP) v112.^3^

**Bioanalytical method for OCT substrate analysis in plasma and urine**

*Plasma:* Metformin, N1-methylnicotinamide (NMN), N1-methyladenosine (NMA), and cimetidine were measured in plasma samples following precipitation with acetonitrile containing an internal standard cocktail (13.3 ng/mL metformin-d6, 10 ng/mL NMN-d3, and 10 ng/mL NMA-d3). The cocktail (400 µL) was added to plasma (100 μL), vortexed for 5 min, and centrifuged at 16,000*g* for 10 min. The supernatant (430 μL) was transferred into a clean tube and dried using a centrifugal concentrator (Concentrator plus/Vacufuge plus, Eppendorf, Germany). The dried sample was reconstituted with 100 µL of acetonitrile:water (5:95, v/v) containing 0.1% formic acid, then vortex mixed for 5 min. After centrifugation at 16,000*g* for 10 min, the supernatant (90 μL) was transferred into a clean tube. The final reconstituted sample was transferred into an LC-MS vial and injected (1 μL) into the LC-MS for quantification of metformin, NMN, NMA, and cimetidine. An exploratory targeted analysis for other potential OCT substrates, including (isobutyryl-L-carnitine) IBC, glycine betaine, tryptophan, N-methyl nicotinic acid, choline, acetylcholine, and carnitine, was conducted using similar processing and analysis methods as described above except that the dried samples were reconstituted with 100 µL acetonitrile:water (30:70, v/v) containing 0.1% formic acid. IBC was quantified in plasma samples using synthetic IBC standard, whereas relative area ratios of the characteristic multiple reaction monitoring (MRM) signals of the remaining potential OCT biomarkers were compared between the two study arms as authentic standards were not available.

For calibration curve preparation, metformin, NMN, NMA, IBC, and cimetidine were serially diluted using methanol:water (80:20, v/v) to produce 10X working concentrations. The working solution (10 µL) was diluted into 2% bovine serum albumin (BSA) matrix (90 μL) to achieve linear calibration curves ranging from 1.3-669.1 ng/mL (metformin), 6.25-200 ng/mL (NMN), 0.4-3.125 ng/mL (NMA), 3.4-216.9 ng/mL (IBC), and 2.0-1022 ng/mL (cimetidine). Low, middle, and high quality control (QC) samples were prepared at concentrations of 9.0, 270, and 629.8 ng/mL (metformin); 8.1, 80.7, and 188 ng/mL (NMN); 0.54, 2.7, and 8.1 ng/mL (NMA); 17.4, 69.4, and 174 ng/mL (IBC); and 13.7, 412.2, and 962 ng/mL (cimetidine), respectively. The calibration curve standards and QC samples (100 µL) were processed and analyzed as described for plasma samples.

*Urine:* Urine samples were thawed and vortexed for 5 min, after which 20 µL were diluted 20-fold into 380 µL of acetonitrile containing the internal standard cocktail (133.4 ng/mL metformin-d6, 100 ng/mL NMN-d3, and 100 ng/mL NMA-d3). After vortexing for 5 min and centrifugation at 16,000*g* for 10 min, the samples were further diluted 5-fold by transferring 200 µL supernatant into a tube containing 800 µL of acetonitrile:water (5:95, v/v) and 0.1% formic acid. Fifty µL of the final diluted sample (100-fold) was transferred into an LC-MS vial, after which 1 µL was injected into the LC-MS/MS system for metformin, NMN, NMA, and cimetidine quantification. For the urine calibration curve, metformin, NMN, NMA, and cimetidine were serially diluted in methanol:water (80:20, v/v) to produce final concentrations ranging from 0.37-743.5 ng/mL (metformin), 1.3-166.7 ng/mL (NMN), 0.08-5.2 ng/mL (NMA), and 0.5-1022 ng/mL (cimetidine). The low, middle, and high QC samples were 8.7, 349.9, and 699.8 ng/mL (metformin); 7.8, 78.4, and 156.9 ng/mL (NMN); 0.3, 2.0, and 7.8 ng/mL (NMA); and 12, 480.9, and 961.9 ng/mL (cimetidine), respectively. Individual calibration standards and QC samples (20 µL) were processed as described for urine samples.

**Untargeted metabolomics analysis of plasma samples**

Considering that the plasma maximum concentration (C_max_) of cimetidine was observed between 1.5 and 3.5 h, 10 µL of processed plasma samples from 1.5, 2, 2.5, 3, and 3.5 h timepoints for 12 subjects in both arms of the study were pooled and used for untargeted metabolomics. The pooled samples (50 µL for each of the 12 subjects from each arm of the study) were vortexed for 5 min and centrifuged at 16,000*g* for 10 min. Twenty µL of the supernatant were transferred into LC-MS vials and injected into a nano-flow-based liquid chromatography-tandem mass spectrometry system (nanoLC-MS/MS, Thermo Scientific™ Q Exactive™ HF). Chromatographic separation of analytes was achieved using a nanoflow LC with PepMap™ RSLC C18 column (75 µm x 25 cm, 2 µm, Thermo Scientific). The mobile phase consisted of 0.1% formic acid in water (A) and 0.1% formic acid in 80% acetonitrile (B), and the following gradient was applied: 0-1 min (0% B), 1 -11 min (10% to 30% B), 11-16 min (30% to 50% B), 16-25 min (50% to 100% B), and 25-40 min (100% B). The mobile phase flow rate was 300 nL/min and the injection volume was 1 µL. MS ionization was achieved using an EASY spray source, and the samples were analyzed in positive ionization and data-independent acquisition (DIA) mode using positive polarity. In addition, the MS was operated in full scan mode with a scan range of *m/z* 70-750, 120,000 resolution, 300 °C capillary temperature, and 150 ms maximum injection time.

**Statistical analysis**

A sample size (n=16) was determined to achieve a power of 0.80 with an alpha value of 0.05 considering the variability in metformin pharmacokinetics and anticipated differences in the pharmacokinetic measures (≥25%). Statistical comparisons of the various measurements of metformin, NMN, NMA, and IBC between the absence and presence of cimetidine, between males and females, and between different genotype groups were made using the paired two-tailed Student’s t-test (GraphPad Prism, San Diego, CA). *p-values* < 0.05 were considered statistically significant. Statistical analysis of the effect of transporter genetic variants on metformin disposition in the baseline arm was conducted using the Kruskal–Wallis test, followed by Dunn’s multiple comparison test (three genotyping groups) or unpaired Student’s t-test followed by the Mann-Whitney Test (two genotyping groups).

**PBPK model development and verification of the cimetidine-metformin interaction**

A middle-out approach was used, where hepatic OCT1 and renal OCT2 intrinsic clearance values for cimetidine (inhibitor) and metformin (substrate, baseline arm) were optimized by inspection of the simulated plasma concentration-time profile against the observed profiles according to acceptance criteria described below. We assumed that cimetidine functions as a competitive inhibitor of metformin transport through OCT1, OCT2, and MATE1/2K. The inhibition constant (K_i_) values for OCT1 and OCT2 were next adjusted to render the model sensitive to cimetidine inhibition. OCT2-mediated uptake was considered rate-limiting by arbitrarily increasing the renal apical efflux clearance of metformin. Biliary clearance (CL_bile_) of metformin was estimated by fitting the model to the reported 4:1 liver:plasma ratio.^4^ Sensitivity analysis was conducted to assess the effect of variable metformin CL_bile_ on hepatic accumulation and plasma concentration.

PBPK modeling was performed using 5 virtual trials (10 participants per trial, age- and sex-matched). The acceptance criteria included 1) maintaining metformin observed concentrations within the 5^th^ and 95^th^ percentiles of simulated concentrations for both baseline and cimetidine exposure arms and 2) maintaining metformin pharmacokinetic parameters within a 2-fold difference between observed and predicted data. Model performance was evaluated by comparing simulated plasma metformin concentrations with multiple sets of observed plasma concentrations from additional cimetidine-metformin studies that were not used for model development. Models were considered acceptable if the observed metformin plasma concentrations from different studies, both at baseline and upon cimetidine exposure, were within the 5^th^ and 95^th^ percentiles of the simulated concentrations.

**RESULTS**

**Bioanalytical method validation**

*Plasma Analysis:*

Precision (% coefficient of variation, %CV) for the low, middle, and high QC samples were 5%, 6%, and 5% for metformin, 6%, 4%, and 4% for NMN, 12%, 10%, and 15% for NMA, and 6%, 9%, and 10% for cimetidine, respectively. Acceptance accuracy criteria for QC samples were 85-115% for metformin and cimetidine and 80-120% for endogenous metabolites (NMN, NMA, and IBC). QC samples were intermittently analyzed during the sample analysis. Seventeen of 18 QCs passed accuracy acceptance criteria with a range of 86-106% for metformin; 17 of 18 QCs passed accuracy acceptance criteria with a range of 80-120% for NMN; 16 of 18 QCs passed accuracy acceptance criteria with a range of 81-112% for NMA; all QCs passed accuracy acceptance criteria with a range of 100-112% for IBC; and 16 of 18 QCs passed accuracy acceptance criteria with a range of 86-112% for cimetidine.

*Urine Analysis:*

The inter-day %CVs for low, middle, and high QC samples were 3%, 3%, 4% for metformin; 4%, 3%, and 3% for NMN; 12%, 3%, and 4% for NMA; and 7%, 1%, and 4% for cimetidine, respectively. Acceptance accuracy criteria for QC samples ranged from 85-115% for metformin and cimetidine and 80-120% for endogenous metabolites (NMN and NMA). Nine QCs were tested for each compound within all assays and all QCs passed the acceptance accuracy criteria. Assay accuracies for QC samples ranged from 87-105% for metformin, 91-105% for NMN, 78-97% for NMA, and 94-108% for cimetidine.

**SUPPLEMENTARY TABLES**

**Table S1.** Inclusion and exclusion criteria for the clinical pharmacokinetic cimetidine-metformin interaction study.

**Inclusion**

- Males and females aged from 18-65 years and healthy
- Not taking any medications (prescription and non-prescription) or dietary/herbal supplements known to alter the pharmacokinetics of metformin or cimetidine.
- Able to understand the informed consent form
- Able to participate in the study (time, transportation, etc.)

**Exclusion**

- Children aged 17 years or less and adults aged 66 years or more
- Any current major illness or chronic illness such as kidney disease, hepatic disease
- Pregnant or nursing
- History of intolerance or allergy to metformin or cimetidine
- Taking concomitant medications, both prescription and non-prescription (including dietary supplements/herbal products) known to alter the pharmacokinetics of metformin or cimetidine

The screening visit consisted of a medical history review, physical exam, blood collection (to obtain complete blood count with differential, electrolytes, liver function tests, serum creatinine), and urine collection (for routine urine analysis and pregnancy tests).

| **Table S2.** Demographic characteristics of the healthy adults who participated in the clinical pharmacokinetic cimetidine-metformin interaction study. | | | | |
| --- | --- | --- | --- | --- |
| **Participant ID** | **Age (years)** | **Sex** | **Race**^a^ | **Ethnicity**^a^ |
| 001 | 28 | Female | White | Not Hispanic or Latino |
| 002 | 58 | Female | White | Not Hispanic or Latino |
| 003 | 39 | Male | White | Unknown / not reported |
| 004 | 27 | Female | Unknown / not reported | Unknown / not reported |
| 005 | 26 | Female | White | Not Hispanic or Latino |
| 006 | 36 | Male | White | Not Hispanic or Latino |
| 007 | 57 | Female | White | Unknown / not reported |
| 008 | 31 | Male | White | Not Hispanic or Latino |
| 009 | 27 | Male | Black or African American | Not Hispanic or Latino |
| 010 | 64 | Female | White | Not Hispanic or Latino |
| 011 | 56 | Male | White | Not Hispanic or Latino |
| 012 | 24 | Female | White | Not Hispanic or Latino |
| 013 | 28 | Female | Asian | Not Hispanic or Latino |
| 014 | 35 | Male | Asian | Not Hispanic or Latino |
| 015 | 30 | Male | More than one race | Not Hispanic or Latino |
| 016 | 50 | Male | White | Not Hispanic or Latino |
| *^a^Self-reported via a standard form* | | | | |

**Table S3.** Liquid chromatography (LC) methods and parameters for metformin, cimetidine, and potential OCT biomarkers.

| Method and Parameters | | |  |
| --- | --- | --- | --- |
| Parameters | | | |
| LC | M-class Waters UPLC | | |
| MS | Waters Xevo TQ-XS | | |
| Column | Acquity UPLC^®^ HSS T3 (1.8 µm, 1x100 mm) | | |
| Mobile phase A | 0.1% Formic acid in water | | |
| Mobile phase B | 0.1% Formic acid in acetonitrile | | |
| Flow rate (µL/min) | 50 | | |
| Column temperature (^o^C) | 40 | | |
| Injection volume (µL) | 1 | | |
| Gradient program (I) | | |  |
| Time (min) | **%A** | **%B** | **Analytes of interest:**   - Metformin - Cimetidine - N1-methylnicotinamide - N1-methyladenosine |
| 0.00 | 95 | 5 |  |
| 1.00 | 95 | 5 |  |
| 2.50 | 45 | 55 |  |
| 4.20 | 32 | 68 |  |
| 4.30 | 10 | 90 |  |
| 5.30 | 10 | 90 |  |
| 5.40 | 95 | 5 |  |
| 7.50 | 95 | 5 |  |
| Gradient program (II) | | |  |
| Time (min) | **%A** | **%B** | **Analytes of interest:**   - Isobutyryl-L-carnitine - Glycine betaine - Tryptophan - N-methyl nicotinic acid - Choline - Acetylcholine - Carnitine |
| 0.00 | 95 | 5 |  |
| 1.00 | 95 | 5 |  |
| 2.50 | 45 | 55 |  |
| 4.20 | 32 | 68 |  |
| 5.20 | 10 | 90 |  |
| 8.00 | 10 | 90 |  |
| 8.50 | 95 | 5 |  |
| 11.50 | 95 | 5 |  |

| **Table S4.** MS transitions, cone voltage, and collision energies of analytes of interest and internal standards used for multiple reaction monitoring (MRM)-based targeted assays. | | | | | |
| --- | --- | --- | --- | --- | --- |
| **MS method (Positive modes)** | | | | | |
| **Compound** | **Retention time (min)** | **Precursor  *m/z*** | **Product  *m/z*** | **Cone voltage** | **Collision energy** |
| Metformin | 1.9 | 130.1 | 60.1 | 25 | 20 |
|  |  | 130.1 | 70.1 | 22 | 17 |
|  |  | 130.1 | 88.1 | 20 | 15 |
| Metformin-d6 | 1.9 | 136.1 | 77.1 | 20 | 20 |
|  |  | 136.1 | 94.1 | 20 | 15 |
| N1-Methylnicotinamide | 1.9 | 137.1 | 94.1 | 25 | 15 |
| N1-Methylnicotinamide-d3 | 1.9 | 140.1 | 97.1 | 25 | 15 |
| Glycine betaine | 1.9 | 118.1 | 72.1 | 20 | 18 |
|  |  | 118.1 | 59.1 | 20 | 18 |
|  |  | 118.1 | 58.1 | 20 | 18 |
| N-Methyl nicotinic acid | 1.9 | 138.1 | 94.1 | 20 | 18 |
| Choline | 1.9 | 104.2 | 94.1 | 20 | 18 |
| Acetylcholine | 1.9 | 146.1 | 87.0 | 20 | 18 |
|  |  | 146.1 | 60.1 | 20 | 18 |
| Carnitine | 1.9 | 162.1 | 103.0 | 20 | 18 |
|  |  | 162.1 | 85.0 | 20 | 18 |
|  |  | 162.1 | 60.1 | 20 | 18 |
| N1-Methyladenosine | 4.3 | 282.1 | 150.1 | 24 | 23 |
| N1-Methyladenosine-d3 | 4.3 | 285.1 | 153.2 | 24 | 23 |
| Cimetidine | 4.4 | 253.1 | 159.1 | 25 | 26 |
|  |  | 253.1 | 95.1 | 25 | 26 |
| Isobutyryl-L-carnitine | 4.3 | 232.2 | 173.0 | 25 | 18 |
|  |  | 232.2 | 85.0 | 25 | 18 |
| Tryptophan | 4.4 | 205.1 | 188.1 | 25 | 18 |
|  |  | 205.1 | 170.1 | 25 | 18 |
|  |  | 205.1 | 159.1 | 25 | 18 |
|  |  | 205.1 | 144.1 | 25 | 18 |
| MS ionization was achieved using an electrospray ionization (ESI) source with positive polarity. | | | | | |

| **Table S5**. Metformin-specific parameters used for metformin PBPK model development. | | |
| --- | --- | --- |
| **Parameter (unit)** | **Value** | **Source** |
| **Physicochemical properties** | | |
| Molecular weight | 129.16 | Simcyp |
| pK_a_ | 11.8 | Simcyp |
| LogP | −1.43 | Simcyp |
| f_u_ | 1 | Simcyp |
| **Absorption (first-order)** | | |
| k_a_ (h^−1^) | 0.27 | Simcyp |
| f_a_ | 0.7 | Simcyp |
| Lag time (h) | 0.29 | Simcyp |
| P_eff_ (10^-4^ cm/s) | 1.226 | Simcyp |
| **Distribution** | | |
| V_ss_ (L/kg) | 0.67381 | Simcyp |
| K_p_ Kidney | 0.98029 | Simcyp |
| K_p_ Liver | 1.0427 | Simcyp |
| K_p_ Gut | 0.89169 | Simcyp |
| **Kidney transport kinetics** | | |
| OCT2 CL_int_(μL·min^−1^·10^6^ cells) | 10.2 | Optimized |
| OCT2 RAF/REF | 1 | Simcyp |
| MATE1/MATE2K CL_int_ (μL·min^−1^·10^6^ cells) | 1000 | Optimized |
| MATE1/MATE2K RAF/REF | 0.128 | Simcyp |
| CL_PD,kidney_ (μL·min^−1^·10^6^ cells) | 4.26 × 10^−7^ | Simcyp |
| **Liver transport kinetics** | | |
| OCT1 CL_int_ (μL·min^−1^·10^6^ cells) | 2.0 | Optimized |
| OCT1 RAF/REF | 1.84 | Simcyp |
| CL_bile_(μL·min^−1^·10^6^ cells) | 0.7 | Optimized |
| CL_PD,Liver_ (μL·min^−1^·10^6^ cells) | 5.88 × 10^−5^ | Simcyp |
| pK_a,_ acid dissociation constant; LogP, Log partition coefficient; f_u_, fraction unbound; k_a_, absorption rate constant; f_a_, fraction absorbed; P_eff_, intestinal effective permeability; V_ss_, volume of distribution at steady state; K_p_, partition coefficient; CL_int_, intrinsic clearance; RAF, relative activity factor; REF, relative expression factor; CL_bile_, biliary clearance; CL_PD_, passive diffusion clearance | | |

| **Table S6**. Cimetidine-specific parameters used in the metformin PBPK model. | | |
| --- | --- | --- |
| **Parameter (unit)** | **PBPK Model** | **Source** |
| **Physicochemical properties** | | |
| Molecular weight | 252.34 | Simcyp |
| pKa | 6.9 | Simcyp |
| LogP | 0.48 | Simcyp |
| fu | 0.8 | Simcyp |
| **Absorption (First-order absorption model)** | | |
| ka (h^−1^) | 0.36 | Optimized |
| fa | 0.92 | Simcyp |
| Lag time (h) | 0.15 | Simcyp |
| **Distribution** | | |
| Vss (L/kg) | 0.5886 | Simcyp |
| Kp Kidney | 0.85038 | Simcyp |
| Kp Liver | 0.87601 | Simcyp |
| Kp Gut | 0.84099 | Simcyp |
| **Kidney Transport kinetic** | | |
| OCT2 CLint(μL·min^−1^·10^6^ cells) | 3 | Optimized |
| OCT2 RAF/REF | 1 | Simcyp |
| OCT2 Ki (μM) | 6.9 | Optimized |
| MATE1 Jmax | 135.5 | Simcyp |
| MATE2K Jmax | 216 | Simcyp |
| MATE1 Km | 7.7 | Simcyp |
| MATE2K Km | 18.2 | Simcyp |
| MATE1 RAF/REF | 1 | Simcyp |
| MATE2K RAF/REF | 1 | Simcyp |
| MATE1/MATE2K Ki (μM) | 0.407 | Simcyp |
| CL_PD,kidney_ (μL·min^−1^·10^6^ cells) | 1.58 × 10^−5^ | Simcyp |
| **Liver Transport kinetic** | | |
| OCT1 CLint(μL·min^−1^·10^6^ cells) | 4 | Optimized |
| OCT1 RAF/REF | 1 | Simcyp |
| OCT1 Ki (μM) | 5.2 | Optimized |
| CL_bile_(μL·min^−1^·10^6 cells) | 3 | Optimized |
| CL_PD,Liver_ (μL·min^−1^·10^6^cells) | 0.1 | Simcyp |
| pK_a,_ acid dissociation constant; LogP, Log partition coefficient; f_u_, fraction unbound; k_a_, absorption rate constant; f_a_, fraction absorbed; P_eff_, intestinal effective permeability; V_ss_, volume of distribution at steady state; K_p_, partition coefficient; CL_int_, intrinsic clearance; RAF, relative activity factor; REF, relative expression factor; CL_bile_, biliary clearance; CL_PD_, passive diffusion clearance; Ki, inhibition constant; Km, Michaelis-Menten constant; Jmax, maximum transport rate. | | |

| **Table S7.** Effects of *OCT* and *MATE* genetic variants on the clinical pharmacokinetic cimetidine-metformin interaction. | | | | | | | |
| --- | --- | --- | --- | --- | --- | --- | --- |
| **Variant** | | | | | **% Change following cimetidine treatment** ^a^ **(↑/↓) (Cimetidine exposure/baseline)** | | |
| **Transporter (gene name)** | **rs ID  (transcript position and nucleotide change)** | **Consequence** | **Genotype** | **Number of subjects** | **AUC_(0-24 h)_** | **C_max_** | **CL_r_** |
| *OCT1* (*SLC22A1*) | rs683369  (c.480G>C,  p.Leu160Phe) | Missense | GC | 5 | ns | ns | - |
|  |  |  | CC | 11 | 33% (**↑**) * | 48% (**↑**) ** | - |
|  | rs628031  (c.1222A>G,  p.Met408Val) | Missense | AA | 3 | ns | ns | - |
|  |  |  | AG | 7 | ns | ns | - |
|  |  |  | GG | 6 | ns | 51% (**↑**) * | - |
|  | rs2114790299 (c.1276+9_1276+16del TGGTAAGT) | Intronic splice recognition site | ref/ref | 3 | ns | ns | - |
|  |  |  | ref/del | 7 | ns | ns | - |
|  |  |  | del/del | 6 | ns | 51% (**↑**) * | - |
|  | rs72552763 (c.1260_1262del ATG, p.Met420del) | Inframe deletion | ref/ref | 12 | ns | 30% (**↑**) * | - |
|  |  |  | ref/del | 4 | ns | ns | - |
| *OCT2*  (*SLC22A2*) | rs316003  (c.1506G>A, p.Val502=) | Synonymous | GA | 5 | ns | - | ns |
|  |  |  | AA | 11 | 27% (**↑**) * | - | 9% (**↓**) * |
|  | rs2774230 (c.518+32C>G) | Intronic | CG | 4 | ns | - | ns |
|  |  |  | GG | 12 | 35% (**↑**) * | - | 9 % (**↓**) ** |
|  | rs624249  (c.390G>T, p.Thr130=) | Synonymous | GG | 6 | ns | - | ns |
|  |  |  | GT, TT | 9,1 | ns | - | 8% (**↓**) * |
| *MATE1* (*SLC47A1*) | rs2247436  (c.499-4G>A) | Intronic splice recognition site | GG | 12 | 17% (**↑**) * | - | 8% (**↓**) * |
|  |  |  | GA | 4 | ns | - | ns |
|  | rs2247437  (c.499-12G>C) | Intronic splice recognition site | GG | 12 | 17% (**↑**) * | - | 8% (**↓**) * |
|  |  |  | GC | 4 | ns | - | ns |
|  | rs2252281  (c.-66T>C) | 5’UTR | TT | 9 | 27% (**↑**) ^$^ | - | 8% (**↓**) * |
|  |  |  | TC | 5 | ns | - | 11% (**↓**) * |
| *MATE2K*  (*SLC47A2*) | rs4925042  (c.885C>T, p.Tyr331=) | Synonymous | CC | 8 | ns | - | 9% (**↓**) * |
|  |  |  | CT, TT | 6, 2 | ns | - | ns |
|  | rs12943590  (c.-130C>T) | Upstream | CC | 7 | 33% (**↑**) * | - | 9% (**↓**) * |
|  |  |  | CT, TT | 8,1 | ns | - | ns |
|  | rs4924792  (c.345C>A, p.Gly115=) | Synonymous | CC | 7 | 33% (**↑**) * | - | 9% (**↓**) * |
|  |  |  | CA, AA | 8, 1 | ns | - | ns |
| The Matched annotations from NCBI and EMBL-EBI (MANE) transcripts for the genes of interest: *SLC22A1* [*OCT1*] (NM_003057.3), *SLC22A2* [*OCT2*] (NM_003058.4), *SLC47A1* [*MATE1*] (NM_018242.3), and *SLC47A2* [*MATE2K*] (NM_001099646.3). OCT2 and MATE2K are encoded on the reverse strand. AUC_(0-24 h)_, area under the plasma-concentration time curve from time zero to 24 h; C_max_, maximum plasma concentration from time zero to 24 h; CL_r_, renal clearance which was calculated based on (0-24 h) time interval; 5’UTR, 5’ untranslated region; del, deletion; and ins, insertion. Statistical analysis was conducted using two-tailed paired t-tests: *p-value* >0.05 (ns, not significant % change following cimetidine treatment), < 0.05 (*) and < 0.01 (**) in cimetidine exposure arm compared to baseline arm of each genetic variant. Statistical analysis was conducted for metformin disposition in baseline arm according to genotype using unpaired t-tests followed by Mann-Whitney test: *p-value* < 0.05 ($). *OCT1* c.411+232C>G (rs4709400), c.412-43T>G (rs4646272), c.516-26C>T (rs45584532), c.955-7C>T (rs7762846), and c.1498+43C>T (rs2297374), *MATE1* c.922-158G>A (rs2289669), and *MATE2K* c.841+14G>C (rs12942065) intronic genetic variants were not associated with metformin pharmacokinetics neither significance and/or magnitude of cimetidine-metformin interaction. ^a^ % Change does not represent the effect of genetic variants on metformin pharmacokinetics. Instead, it represents the effect of single nucleotide polymorphisms (SNPs) on drug-drug interaction potential. | | | | | | | |

**SUPPLEMENTARY FIGURES**

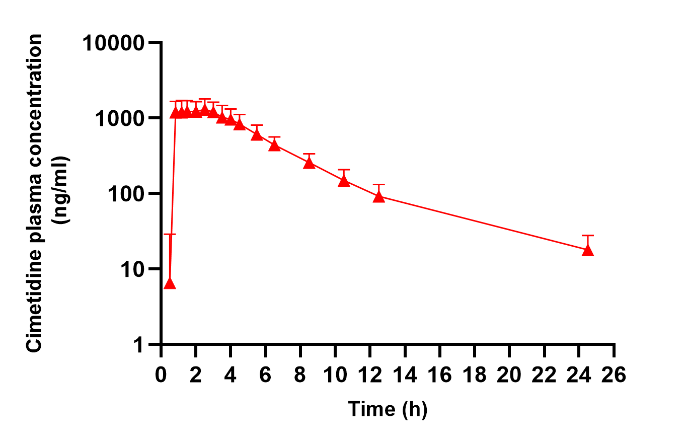

**Figure S1.** Geometric mean plasma concentration-time profile of cimetidine after a single oral dose of 400 mg in healthy subjects (n=16). Symbols and error bars represent observed geometric means and 90% confidence intervals, respectively.

**D**

**B**

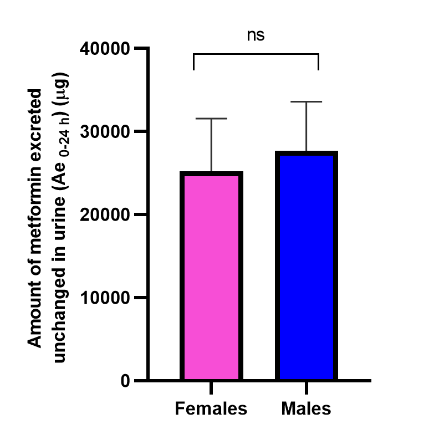

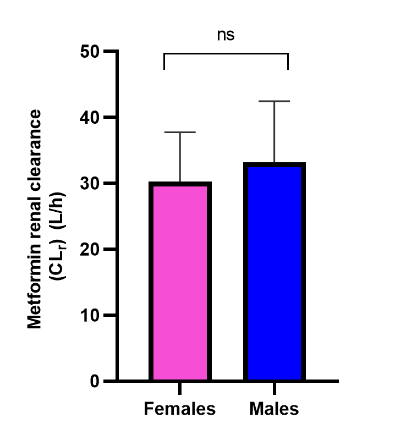

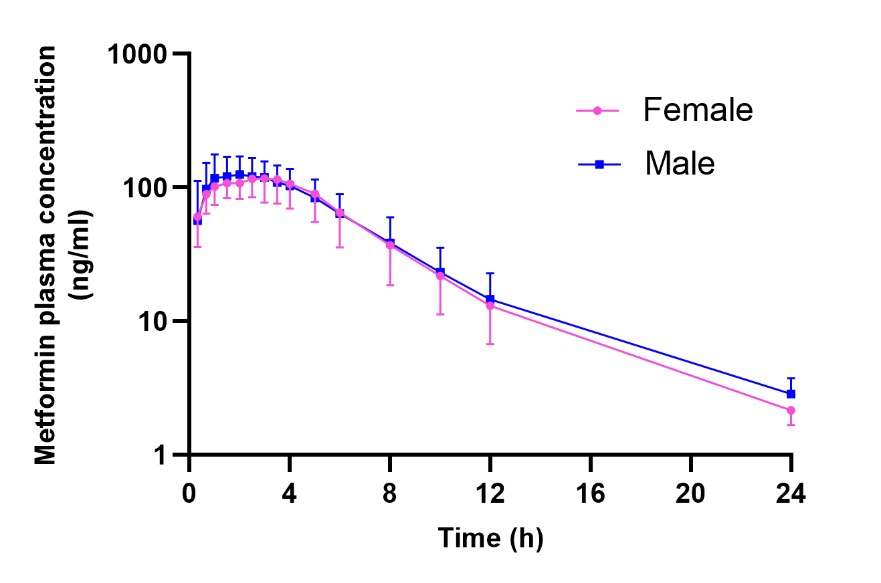

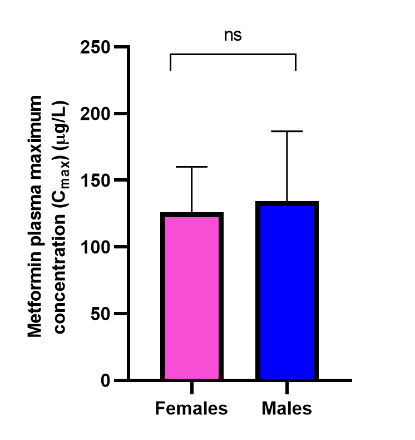

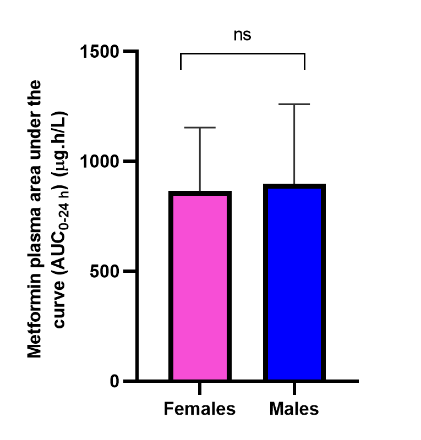

**Figure S2.** Association of sex on the mean plasma concentration-time profile (A), plasma area under the curve from 0-24 h (AUC_0-24 h_) (B), plasma maximum concentration (C_max_) (C), amount excreted into the urine from 0-24 h (Ae_0-24 h_) (D), and renal clearance (CL_r_) (E) of metformin between healthy female (pink) and male (blue) (n=16, 8 females, 8 males) in metformin alone arm (baseline). One male was excluded from the Ae_0-24 h_ and CL_r_ analysis due to difficulty voiding urine. In Figure S2E, CL_r_ was calculated based on (0-24 h) time interval. Symbols and error bars denote observed means and 90% confidence intervals, respectively. Statistical analysis was conducted using two-tailed paired t-tests: *p-value* > 0.05 (ns, not significant).

**E**

**A**

**C**

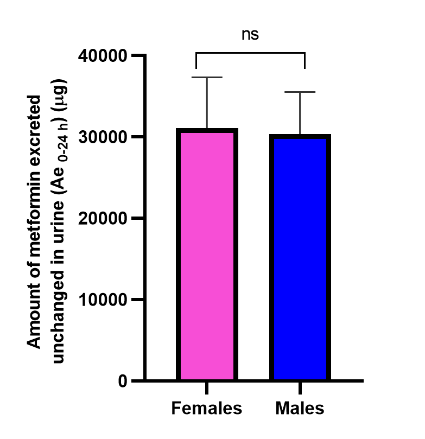

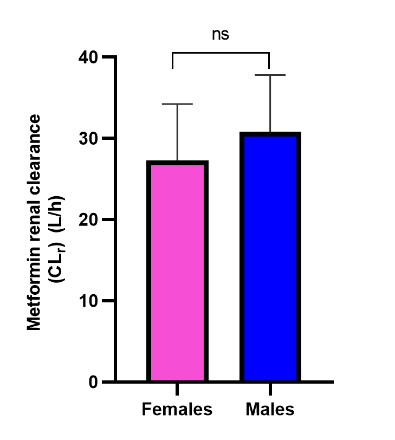

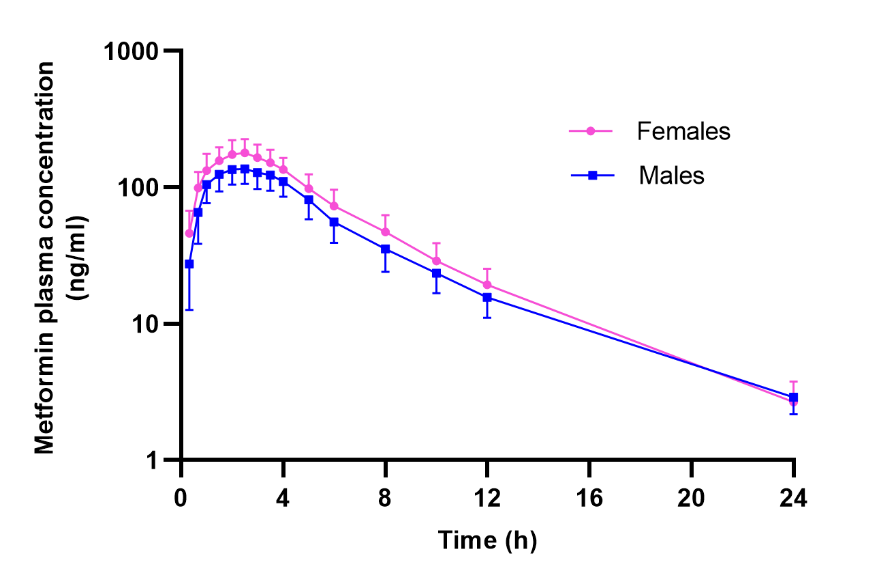

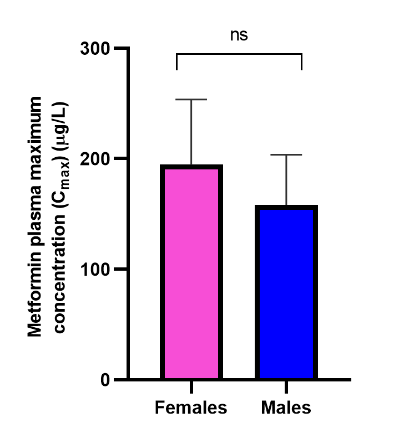

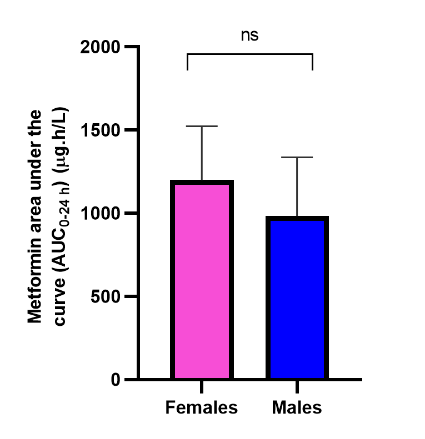

**Figure S3**. Association of sex with mean plasma concentration-time profile (A), plasma area under the curve from 0-24 h (AUC_0-24 h_) (B), plasma maximum concentration (C_max_) (C), amount excreted into the urine from 0-24 h (Ae_0-24 h_) (D), and renal clearance (CL_r_) (E) of metformin between healthy female (pink) and male (blue) (n=16, 8 females, 8 males) in metformin plus cimetidine arm (cimetidine exposure). One male was excluded from the Ae_0-24 h_ and CL_r_ analysis due to difficulty voiding urine. In Figure S3E, CL_r_ was calculated based on (0-24 h) time interval. Symbols and error bars denote observed means and 90% confidence intervals, respectively. Statistical analysis was conducted using two-tailed paired t-tests: *p-value* > 0.05 (ns, not significant).

**D**

**E**

**A**

**C**

**B**

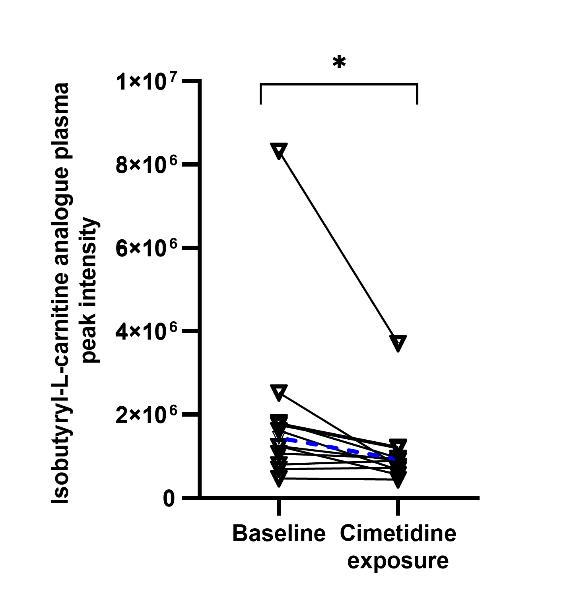

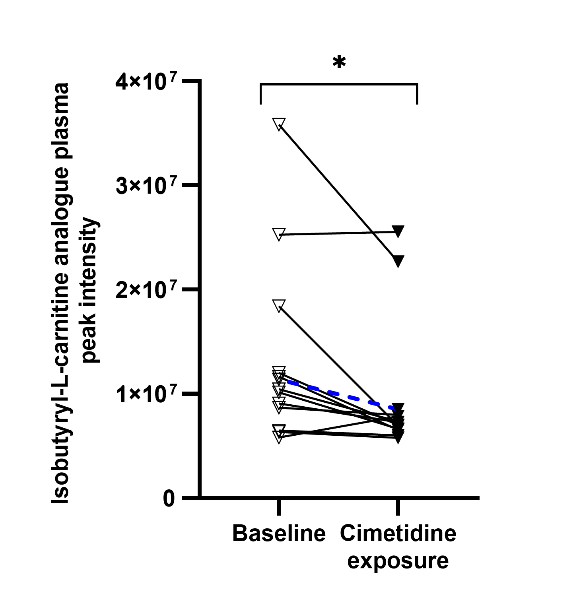

**Figure S4.** Effect of cimetidine co-administration on the known potential endogenous OCT1 biomarker isobutyryl-L-carnitine (IBC) analogues. LC-MS peak areas of IBC analogues; *m/z* 232.1541 with retention time 22.3 (A) and *m/z* 232.1542 with retention time 23.5 (B) in pooled plasma (n=12) at 1.5, 2, 2.5, 3, and 3.5 h in the metformin alone (baseline, open symbols) and metformin plus cimetidine (cimetidine exposure, solid symbols) arms among healthy adults. Solid lines indicate individual values and dashed line represents geometric mean. Statistical analysis was conducted using two-tailed paired t-tests: *p-value* < 0.05 (*).

**B**

**A**

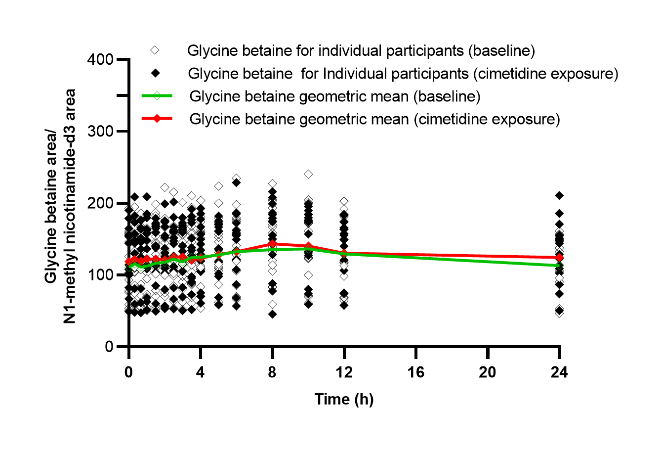

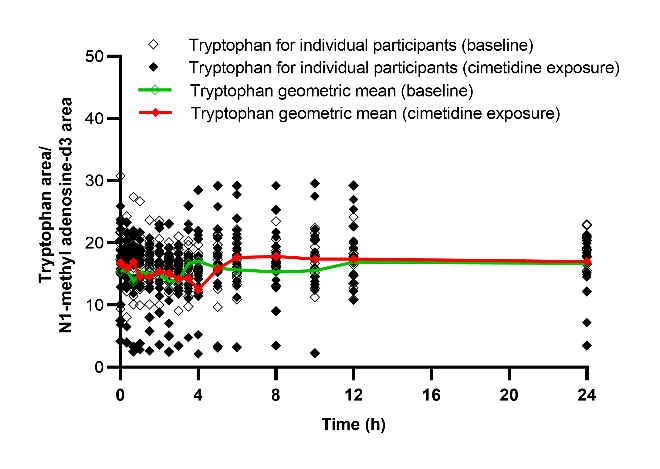

**A**

**B**

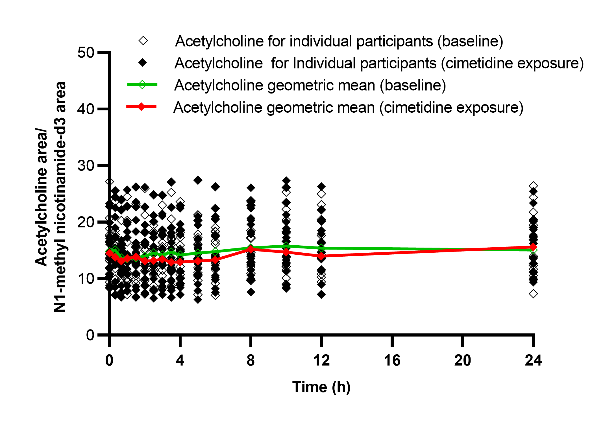

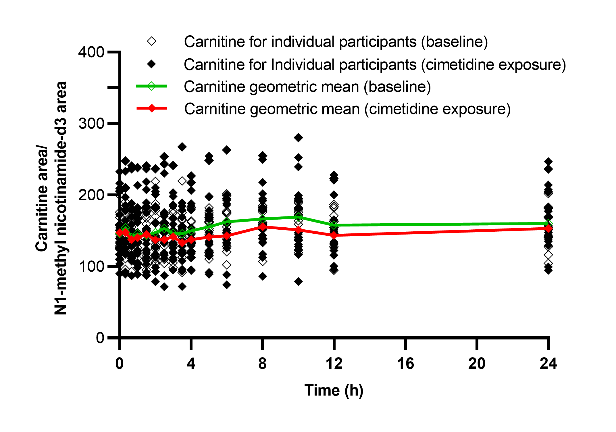

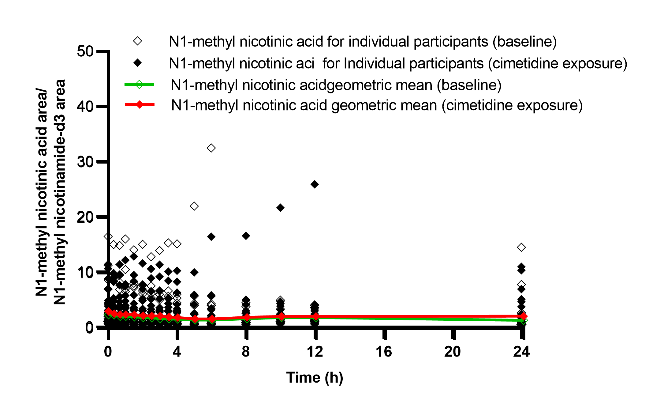

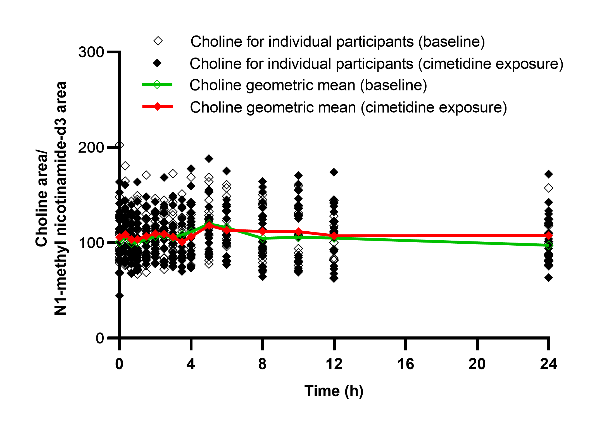

**Figure S5.** Effect of cimetidine co-administration on the potential OCT substrates. LC-MS/MS area ratio of glycine betaine (A), tryptophan (B), N1-methyl nicotinic acid (C), choline (D), acetylcholine (E), and carnitine (F) in the metformin alone (baseline, open symbols) and metformin plus cimetidine (cimetidine exposure, solid symbols) arms among healthy adults (n=16).

**E**

**F**

**D**

**C**

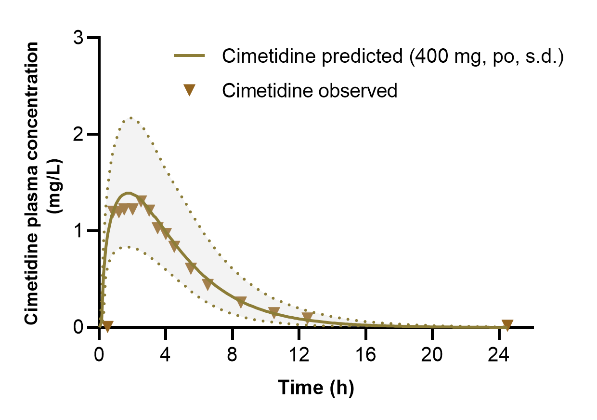

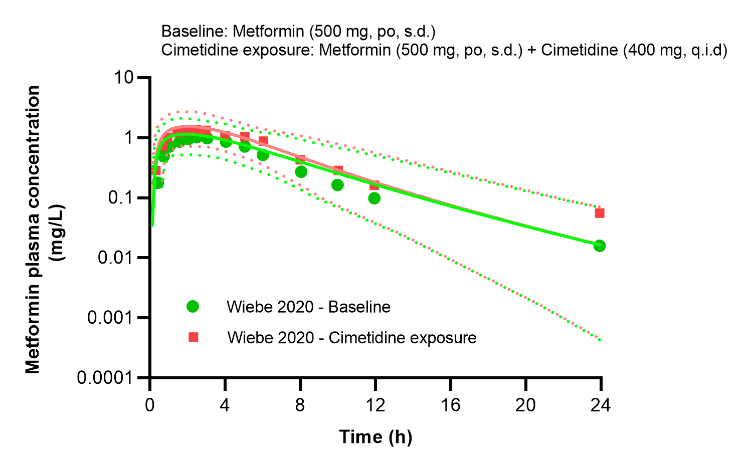

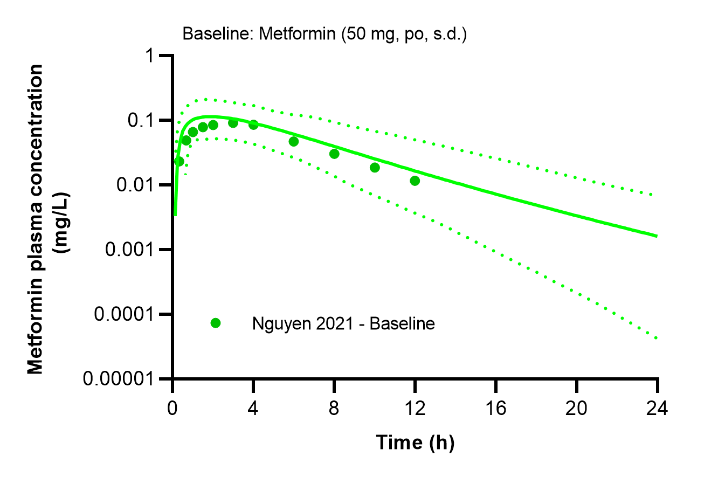

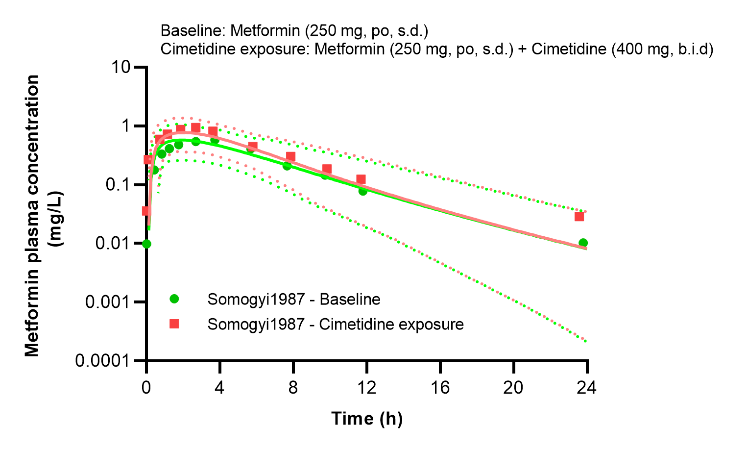

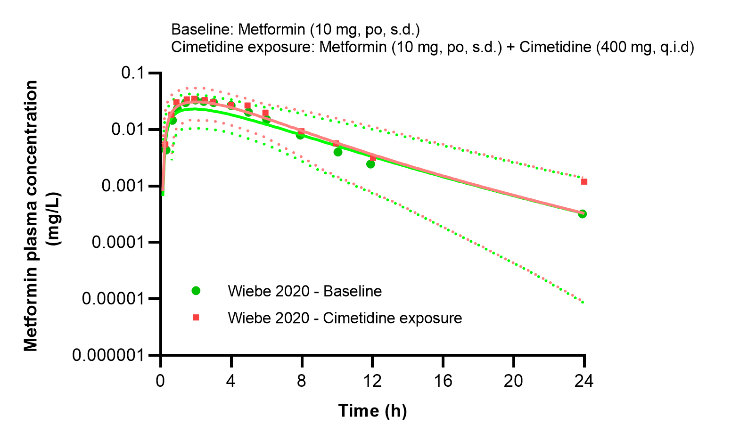

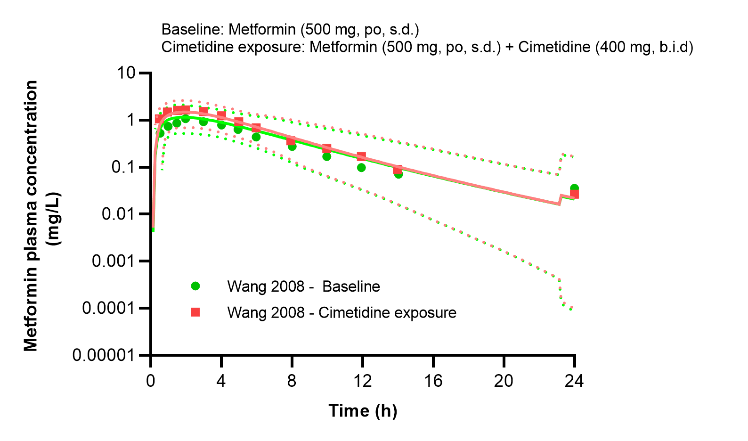

**Figure S6.** Physiologically based pharmacokinetic (PBPK) model predicted mean cimetidine plasma concentration-time profiles from our study (A) after 400 mg single dose of cimetidine. Metformin PBPK model for metformin subtherapeutic dose (50 mg) was reproduced from Nguyen et al. 2021 (B). Cimetidine-metformin PBPK model predicted mean metformin plasma concentration for metformin at 10 (C), 250 (D), and 500 mg (E, F) doses. Population prediction arithmetic means of metformin plasma concentration are shown as solid green (baseline) and solid red (cimetidine exposure) lines. 5^th^ and 95^th^ percentiles of the predicted metformin plasma concentrations are illustrated with the area constrained by green (baseline) and red (cimetidine exposure) dash lines. The observed data are shown as green dots (baseline) and red squares (cimetidine exposure).

**A**

**F**

**B**

**D**

**E**

**C**

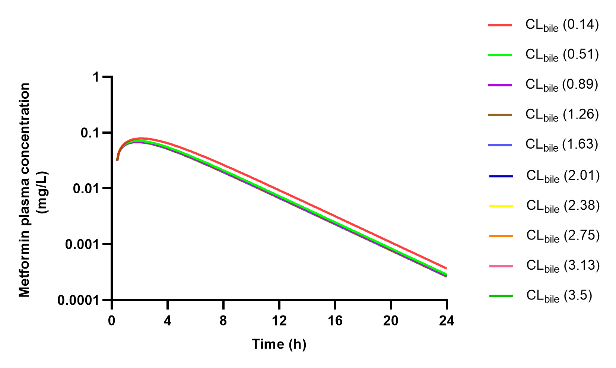

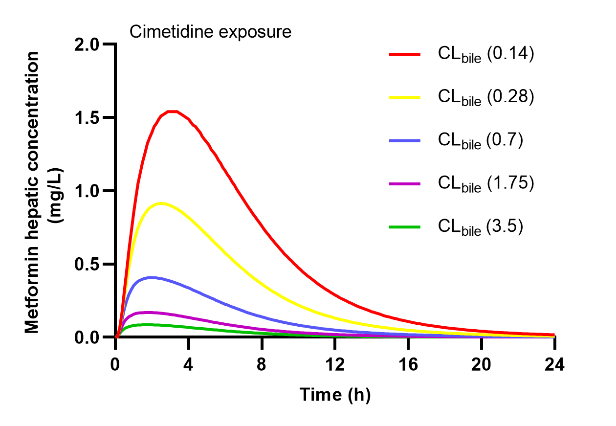

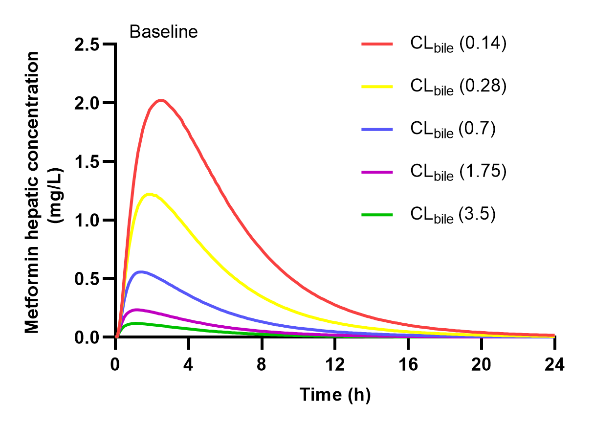

**C**

**A**

**Figure S7.** Local sensitivity analysis of biliary secretion (CL_bile_) in metformin physiologically based pharmacokinetic (PBPK) model. The PBPK model predicted impact of CL_bile_ on metformin hepatic concentration when administered alone (baseline) (A) and in combination with cimetidine (cimetidine exposure) (B), and metformin plasma concentration (C).

**B**
